# Alcoholism Gender Differences in Brain Responsivity to Emotional Stimuli

**DOI:** 10.1101/428565

**Authors:** Kayle S. Sawyer, Nasim Maleki, Trinity Urban, Ksenija Marinkovic, Steven A. Karson, Susan M. Ruiz, Gordon J. Harris, Marlene Oscar-Berman

## Abstract

Men and women may use alcohol to regulate their emotions differently, with corresponding differences in neural responses. We examined how photographs of emotional stimuli impacted brain activity obtained through functional magnetic resonance imaging (fMRI) from 42 alcoholic (25 women) and 46 nonalcoholic (24 women) participants. Brain responsivity was blunted in alcoholic compared to nonalcoholic groups. Further analyses indicated significant gender differences in the impact of alcoholism. Brain activation of the alcoholic men (ALC_M_) was significantly lower than that of the alcoholic women (ALC_W_) and nonalcoholic men (NC_M_) in regions including the inferior parietal gyrus, anterior cingulate gyrus, and postcentral gyrus, whereas activation was higher in the ALC_W_ than in the nonalcoholic women (NC_W_) in superior frontal and supramarginal cortical regions. The reduced brain reactivity of ALC_M_ and increases for ALC_W_ highlighted divergent brain regions and gender effects, suggesting possible differences in the underlying basis for development of alcohol use disorders.

## Introduction

Impaired affect regulation is a primary motive for the use of drugs, including alcohol (Prescott, Cross, Kuhn, Horn, & Kendler, 2004; Vaughan et al., 2012). Affective processing deficits have been linked to misinterpretation of environmental cues, irregularity in mood, and increased alcohol consumption and may be an underlying factor leading to the development and maintenance of alcohol use disorders (AUD) (Gilman, Ramchandani, Davis, Bjork, & Hommer, 2008; Thorberg, Young, Sullivan, & Lyvers, 2009). However, problem drinkers are a heterogeneous population. While alcohol and other GABAergic agents such as benzodiazepines typically are considered to be depressants because of their ability to decrease anxiety, tension, and inhibition, they also can function as a stimulant, generating feelings of euphoria and well-being (Gilman et al., 2008; Mukheijee, Das, Vaidyanathan, & Vasudevan, 2008). This may make alcohol appealing to men and women subgroups of problem drinkers for contrasting reasons (Buchmann et al., 2010). For example, on average, women might drink to decrease aversive affect, and men might drink to enhance favorable emotional states (Buchmann et al., 2010; Buckner, Eggleston, & Schmidt, 2006; Crutzen, Kuntsche, & Schelleman-Offermans, 2013; Oscar-Berman et al., 2014; S. Ruiz & Oscar-Berman, 2015).

Research has indicated that men and women process emotions differently (Mareckova et al., 2016; Proverbio, Adorni, Zani, & Trestianu, 2009). Women on average display different psychophysiological reactions to emotional stimuli (Sawyer, Poey, Ruiz, Marinkovic, & Oscar-Berman, 2015) and tend to be more emotionally expressive than men (Kring & Gordon, 1998). Conversely, men on average have an increased tendency to repress emotional responses (Birditt & Fingerman, 2003). Additionally, alcoholic women are two to three times more likely to be diagnosed with anxiety and affective disorders than alcoholic men, while ALC_M_ are twice as likely as ALC_W_ to have antisocial personality disorders (Merikangas et al., 1996; Oscar-Berman et al., 2009). The presence of gender-specific deficits in emotional regulation may provide insight into what differentially motivates men and women to abuse alcohol (Erol & Karpyak, 2015; Mosher Ruiz, Oscar-Berman, Kemppainen, Valmas, & Sawyer, 2017; Regier et al., 1990; S. Ruiz & Oscar-Berman, 2015; Valmas, Mosher Ruiz, Gansler, Sawyer, & Oscar-Berman, 2014).

The presentation of emotional stimuli is associated with activity within a well-characterized neural system encompassing the prefrontal cortex, insula, cingulate cortex, and medial temporal lobe structures including the amygdala (R. J. Davidson, Abercrombie, Nitschke, & Putnam, 1999; Proverbio et al., 2009). Alcoholics exhibit reduced fMRI activation in the amygdala, hippocampus, and anterior cingulate cortex while viewing emotional faces, compared to nonalcoholic control participants (Marinkovic et al., 2009; Salloum et al., 2007). This reduced emotional responsivity could reflect deficits in the accurate interpretation and communication of emotional states, contributing to personality disorders and social impairments that characterize alcoholics (Oscar-Berman et al., 2009, 2014). However, prior research in this area traditionally has considered only male alcoholics or has collapsed participants across both genders. Whereas brain activation alterations in emotional processing have been studied in relation to AUD (Beck et al., 2009; Chanraud-Guillermo et al., 2009; Gilman & Hommer, 2008; Heinz et al., 2007), gender differences have not been explored in depth, and there is a need for more research in this domain (Nixon, Prather, & Lewis, 2014; S. M. Ruiz & Oscar-Berman, 2013). In any case, a more thorough understanding of the associations among gender, emotional reactivity, and alcohol consumption could potentially allow for the development of individually tailored treatments.

In the present study, we sought to characterize gender differences in the relationship between long-term alcoholism and the neural networks that support emotion perception. We expected that alcoholic participants (ALC) would exhibit blunted emotional reactivity compared to nonalcoholic controls (NC) during the processing of emotional stimuli, and that these relationships would indicate a different pattern of abnormalities for men and women.

## Methods

### Participants

Prior to conducting the experiment, we computed estimates of sample size based upon Cohen’s *d*, which suggested approximately 20 participants per group were required to detect a medium to large effect size (Cohen, 1988), a number confirmed by fMRI-specific research (Thirion et al., 2007). A total of 88 participants (25 ALC_W_, 17 ALC_M_, 24 NC_W_, and 22 NC_M_) were included in the analyses. The characteristics of the participants, including alcoholism indices and neuropsychological test scores are presented in Table 1 of the Results section; data and code are available from DRYAD and GitLab (https://gitlab.com/kslavs/sawver-iaps). All participants were right-handed English speakers recruited through flyers placed in facilities and in public places (e.g., churches, stores), and advertisements placed with local newspapers and websites. Selection procedures included an initial structured telephone interview to determine age, level of education, health history, and history of alcohol and drug use. Eligible individuals were invited to the laboratory for further screening and evaluations ranging between five to eight hours over the course of one to three days. Prior to screening, informed consent was obtained; the protocols and consent forms were approved by the Institutional Review Boards of the participating institutions. Participants were reimbursed for their time and travel expenses.

Participants underwent medical history interview and vision testing, plus a series of questionnaires (e.g., handedness, alcohol and drug use, Hamilton Rating Scale for Depression (HRSD; (Hamilton, 1960)) to ensure they met inclusion criteria. Participants were given the computerized Diagnostic Interview Schedule (Robins et al., 2000), which provides lifetime psychiatric diagnoses according to criteria established by the American Psychiatric Association. Participants were excluded from further participation if any source (e.g., hospital records, referrals, or personal interviews) indicated that they had one of the following: Corrected visual acuity worse than 20/50 in both eyes; Korsakoff’s syndrome; cirrhosis, major head injury with loss of consciousness greater than 15 minutes unrelated to AUD; stroke; epilepsy or seizures unrelated to AUD; schizophrenia; HRSD score over 15; electroconvulsive therapy; history of illicit drug use more than once per week within the past five years (except for one ALC_W_ who had used marijuana more frequently but not during the six months preceding testing, and one ALC_W_ who had used marijuana once per week for four years, ceasing four years before testing); lifetime history of illicit drug use more than once per week for over 10 years or three times per week for over five years.

Participants received a structured interview regarding their drinking patterns, including length of abstinence and duration of heavy drinking, i.e., more than 21 drinks per week (one drink: 355 ml beer, 148 ml wine, or 44 ml hard liquor). For each participant, we calculated a Quantity Frequency Index (Cahalan, Cisin, & Crossley, 1969), which factors the amount, type, and frequency of alcohol usage (ounces of ethanol per day, roughly corresponding to number of drinks per day) over the last six months (for the NC group), or over the six months preceding cessation of drinking (for the ALC group). The ALC participants met criteria for alcohol abuse or dependence, and had over 21 drinks per week for at least five years in their lifetime; all had abstained from alcohol for at least 21 days. None of the NC participants drank heavily (21 or more per week), except for one man who drank while serving in the army decades before the scan, but did not meet the criteria for alcohol dependence. We examined the group x gender interaction within a regression model for the demographics, alcoholism indices, neuropsychological and clinical assessment scores. We also conducted Welch’s t-tests to examine gender differences for each measure for the ALC and NC groups separately, and group differences for the men and women separately.

### MRI Acquisition

Imaging data were acquired using a 3T Siemens (Erlangen, Germany) Trio Tim magnetic resonance scanner. Following automated shimming and scout image acquisition, two eight-minute 3D Tl-weighted MP-RAGE sequences were obtained: TR = 2530 msec, TE = 3.45 msec, flip angle = 7^0^, FOV = 256 mm, 128 sagittal slices with in-plane resolution 1 × 1 mm, slice thickness = 1.33 mm. These two structural volumes were used for functional slice prescription, spatial normalization, and cortical surface reconstruction. Due to time constraints, only one MP-RAGE sequence was obtained for 23 subjects (11 NC_M_, 8 ALC_M_, 2 NC_W_, 2 ALC_W_). Functional whole-brain BOLD (blood oxygen level-dependent) images were obtained with a gradient echo T2*-weighted sequence: TR = 2 sec, TE = 30 msec, flip angle = 90°, FOV = 200 mm, slice thickness = 3.0 mm, spacing =1.0 mm, 32 interleaved axial-oblique slices aligned to the anterior-commissure/posterior-commissure line (voxel size: 3.1 × 3.1 × 4.0 mm). The scans covered the entire cerebrum and the superior portion of the cerebellum.

### Behavioral Task

Participants were presented with blocks of pictures chosen to evoke emotional responses (**Figure 1**). The picture stimuli were from the International Affective Picture System (Lang, Ohman, & Vaitl, 1988). Participants completed five runs (except one NC_W_ who completed only four runs), each including five conditions: aversive, erotic, gruesome, happy, and neutral pictures. As depicted in Figure 1, each run contained three 24-second blocks of fixation plus eight 24-second blocks that each consisted of six pictures of one of the emotional conditions (e.g., happy pictures), for a total of 11 blocks per run. The five runs included a total of 40 blocks of emotional pictures with eight blocks for each of the five emotional picture conditions. Stimuli were presented only once, totaling 48 pictures per 264-second run (240 pictures in 22 minutes in total across the five runs).

Within stimulus blocks, the six pictures were each serially presented against a black background for 3 seconds, followed by 1 second of fixation (+++). Participants were instructed to answer the question: “How does the picture make you feel?” Following each image within a block, participants indicated feeling *good*, *neutral*, or *bad*, by using their index fingers to press buttons on a box. The left index finger indicated *good*, the right index finger indicated *bad*, and both center buttons indicated *neutral*; the left and right were counterbalanced across participants. Block order was counterbalanced across runs, and run order was counterbalanced across participants. The task was presented with the Presentation software package (Neurobehavioral Systems, Albany, CA, USA).

**Figure 1.**
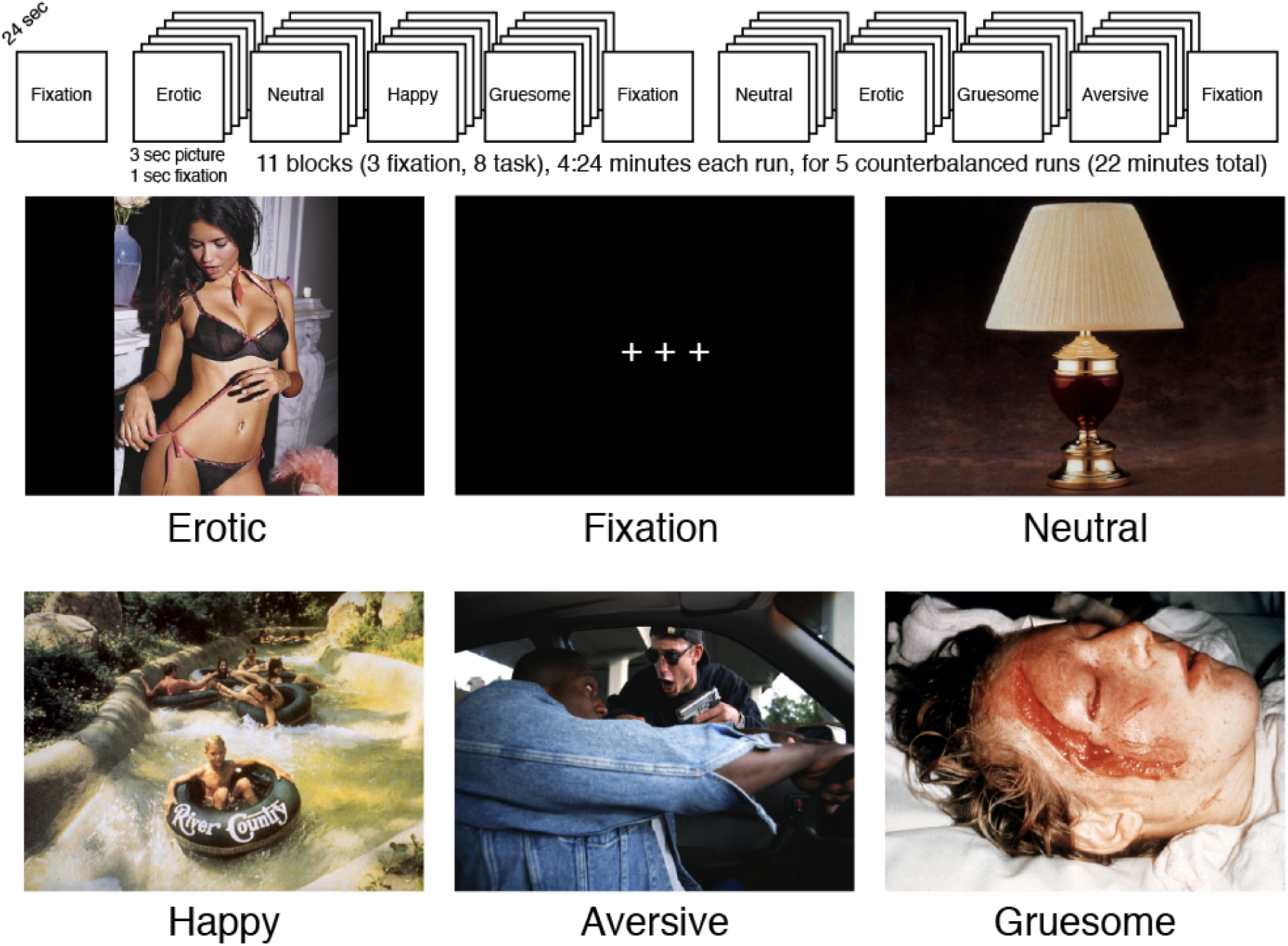
Schematic of task presentation, and examples of stimuli. As described in the text, participants were shown pictures from the IAPS and asked to report how the pictures made them feel (*good*, *bad*, or *neutral*).

Behavioral response data were analyzed using R software mixed models (Bates, Maechler, Bolker, & Walker, 2018; R Core Team, 2017), with one model specified for reaction times, and one model specified for the percentage of pictures endorsed for each rating (good, neutral, bad). For both reaction times and percentage models, independent intercepts were modeled for each participant, and full-factorial ANOVAs were calculated for the four factors of rating (good, neutral, bad), condition (aversive, erotic, gruesome, happy, neutral), group (ALC, NC), and gender (men, women).

### MR1 Analyses

The imaging data were analyzed using FreeSurfer and FS-FAST v6.0 (http://surfer.nmr.mgh.harvard.edu) analysis packages (Dale, Fischl, & Sereno, 1999; B. Fischl, Sereno, & Dale, 1999). Individual cortical surfaces were reconstructed using automatic gray and white matter segmentation, tessellation, and inflation. Images were registered with a canonical brain surface (fsaverage) based on sulcal and gyral patterns (B. Fischl, Sereno, Tootell, & Dale, 1999), and registered with a canonical brain volume (MNI305) using a 12 degrees of freedom nonlinear transform. Gray and white matter surface accuracy were individually examined using automatically-generated quality control figures (https://github.com/poldracklab/niworkflows), and no errors were detected that would be likely to influence the outcomes of this project (Waters, Mace, Sawyer, & Gansler, 2018).

The fMRI data were corrected for motion and slice-time acquisition using FS-FAST preprocessing. Normalized motion and signal intensity spikes were obtained from the nypipe rapidart algorithm (https://www.nitre.org/proiects/rapidart/, https://doi.org/10.5281/zenodo.596855), and blocks with motion over 1.5 mm, or signal intensity shifts over 3.0 standard deviations, were removed via a paradigm file covariate for each run. Subjects were removed from the study if this process excluded all but three or fewer blocks of any condition. Next, the FS-FAST process split the analysis into three spaces (left and right surfaces, and subcortical volume), then data from each subject was spatially normalized (co-registered with) the fsaverage and MNI305 spaces, respectively; all subsequent analyses were performed in these three group spaces. Spatial smoothing was performed with a 5 mm full width at half maximum Gaussian kernel in 3D for the volume and in 2D for the surfaces. Condition-specific effects were estimated by fitting the amplitudes of boxcar functions convolved with the FSL canonical hemodynamic response function to the BOLD signal across all runs.

Statistical maps were constructed from each contrast of stimulus conditions for each subject (first level analyses). Four contrasts were examined: aversive vs. neutral, happy vs. neutral, erotic vs. neutral, and gruesome vs. neutral; and an additional contrast of neutral vs. fixation was performed as context. These first-level analyses were concatenated, and second-level (group level or between-subjects) analyses were performed using random-effects models to account for inter-subject variance (Friston, Holmes, Price, Büchel, & Worsley, 1999), with weighted least squares effects incorporated from the variability measures from the first-level contrasts. We examined the overall main effect of group (ALC vs. NC), the interaction of group x gender, and the effects of group for men and women separately, for each of the four contrasts (each emotion condition vs. neutral condition), along with the neutral vs. fixation contrast. For the cortical surface maps, cluster-level corrections for multiple comparisons were applied to each map (Hagler, Saygin, & Sereno, 2006) using 10,000 precomputed Z Monte Carlo simulations with two threshold criteria: *p<*0.05 at each vertex, and *p*<0.05 for the entire cluster (further corrected for the analysis of three spaces: left cortex, right cortex, and subcortical). Cortical surface cluster regions were identified by the location of each cluster’s peak vertex on the cortical surface (Desikan et al., 2006), and subcortical cluster regions were identified by the MNI coordinates of each cluster’s peak voxel (Bruce Fischl et al., 2002).

## Results

### Participant Characteristics

Demographics, alcoholism indices, neuropsychological and clinical assessment scores of the 88 participants are presented in **Table 1**. Although the HRSD scores for the ALC men and women were higher than for the NC men and women (*p*<0.01), both groups’ scores were very low (mean 3.6 vs. 1.1): HRSD scores of 8, 16, and 25 or above indicate mild, moderate, or severe depression, respectively (Zimmerman, Martinez, Young, Chelminski, & Dalrymple, 2013). The average number of daily drinks (DD) was significantly higher in ALC_M_ compared to ALC_W_ (*p*<0.05), Both alcoholic and control women had higher delayed memory scores compared to men in their respective group (Wechsler Memory Scale Delayed (General) Memory Index, *p*<0.01).

**Table 1.** Participants’ characteristics and drinking measures. Values presented as mean ± standard deviation. Abbreviations: ALC_W_ = Alcoholic women; ALC_M_ = Alcoholic men; NC_W_ = Nonalcoholic control women; NC_M_ = Nonalcoholic control men; DHD = Duration of Heavy Drinking (>21 drinks per week) in years; DD = Daily drinks; LOS = Length of sobriety in years. HRSD = Hamilton Rating Scale for Depression (Hamilton, 1960); WMS DMI = Wechsler Memory Scale, 3rd ed. Delayed (General) Memory Index; NA = Not Applicable. Significant differences: ^a^(ALC_M_ > NC_M_, *p*<0,05): ^b^(ALC_M_ > NC_M_, *p<*0.001): ^c^(ALC_W_ > NC_W_, *p*<0.01); ^d^(ALC_w_ > NC_W_, *p*<0.001); ^e^(ALC_w_ > ALC_M_, *p*<0.001); ^f^(ALC_M_ > ALC_w_, *p*<0.05); ^g^(NC_W_ > NC_M_, *p*<0.05); ^h^(NC_w_ > NC_m_, *p*< 0.01); ‘(group x gender interaction, *p<*0.05).

| Measure | ALC<sub>W</sub><br>N = 25 | ALC<sub>M</sub><br>N = 17 | NC<sub>W</sub><br>N = 24 | NC<sub>M</sub><br>N = 22 |
| --- | --- | --- | --- | --- |
| Age | 52.0 ± 10.6 | 53.2 ± 9.7 | 54.4 ± 15.4 | 55.0 ± 12.4 |
| Education<sup>g</sup> | 15.3 ± 2.3 | 13.8 ± 2.5 | 16.1 ± 2.6 | 14.8 ± 1.9 |
| VIQ | 110.4 ± 16.6 | 107.0 ± 15.0 | 113.2 ± 17.8 | 109.9 ± 11.1 |
| PIQ | 106.9 ± 17.7 | 100.1 ± 12.5 | 111.2 ± 16.9 | 107.1 ± 11.8 |
| WMS DMI<sup>e,h</sup> | 119.1 ± 15.9 | 99.0 ± 10.3 | 117 ± 17.3 | 105.0 ± 14.0 |
| HRSD<sup>a,c</sup> | 3.4 ± 3.5 | 3.6 ± 4.7 | 1.2 ± 2.2 | 1.0 ± 1.2 |
| DHD<sup>b,d</sup> | 13.3 ± 6.4 | 14.6 ± 6.2 | 0.0 ± 0.0 | 0.0 ± 0.0 |
| DD<sup>b,d,f,i</sup> | 6.9 ± 6.3 | 12.9 ± 9.6 | 0.2 ± 0.3 | 0.4 ± 0.5 |
| LOS | 7.3 ± 8.9 | 7.5 ± 11.9 | NA ± NA | NA ± NA |

### Behavioral Ratings

Of the 88 participants included in fMRI analyses, 12 were excluded from the analysis of behavioral ratings because of technical problems or incomplete data, leaving 76 subjects for the final analyses (21 ALC_W_, 15 ALC_M_, 21 NC_W_, 19 NC_M_). Overall, participants percentage ratings of *good*, *bad*, and *neutral* were generally consistent among ALC and NC men and women (**Figure 2**) for the various conditions (aversive, erotic, gruesome, happy, neutral). That is, the participants rated erotic pictures as mostly *neutral* and *good*; gruesome pictures as almost entirely *bad*; aversive pictures as *bad*, with a few *neutral*; happy pictures as mostly *good*, with some *neutral*, and neutral pictures as mostly *neutral*, with some *good* (altogether representing a significant condition x rating interaction, **Figure 2-S1**). While all groups (ALC and NC men and women) had a similar pattern, a significant group x condition x rating interaction revealed that the ALC group rated erotic pictures as *good* less frequently than the NC group. The gender x condition x rating interaction revealed that more men than women rated erotic pictures as *good*.

**Figure 2.**
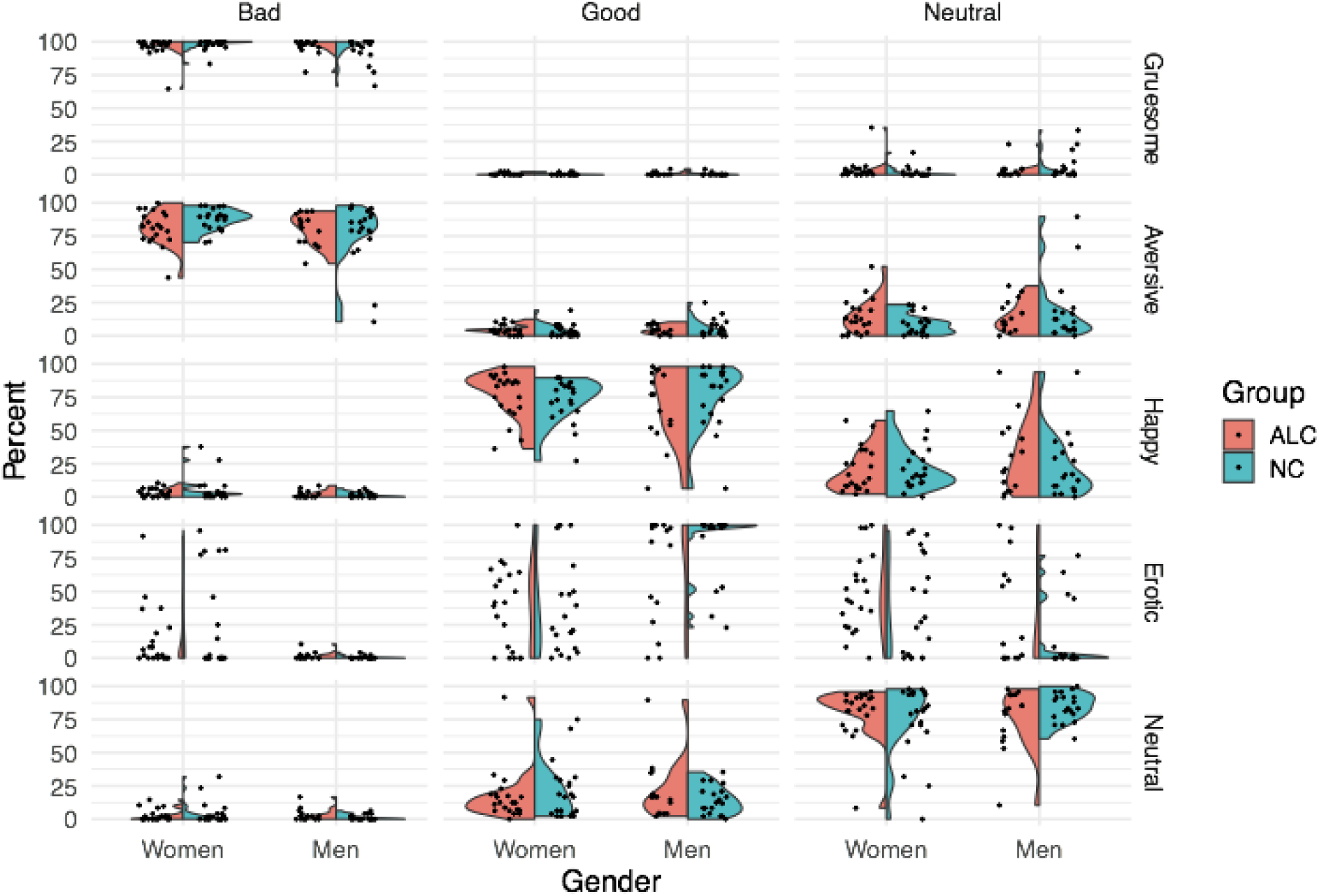
Percentage of behavioral ratings by condition, rating, group, and gender. The split violin plot represents the significant condition x rating x group interaction, and the significant condition x rating x gender interaction, for percentage rating of the pictures *p*<0,05 (Figure 2-S1). The group interaction is most clearly evident for the difference in the *good* and *neutral* ratings of the erotic pictures, with the alcoholic participants rating the pictures *good* less frequently; other picture types were rated more similarly by both the alcoholic and control groups. The gender interaction indicated that men rated erotic pictures as *good* more frequently than women. Figure 2-S2 shows the reaction times. Abbreviations: ALC = Alcoholic participants; NC = Nonalcoholic Control participants.

As with the percentage ratings, evaluation times also were generally comparable for the four groups (**Figure 2-S2** and **Figure 2-S3**). There were significant interaction effects of condition x rating, rating x gender, and main effects of condition and rating (*p*<0.001 for all). The evaluation time for gruesome and aversive stimuli were approximately 0.5 seconds longer than other conditions. The evaluation time for *bad* ratings were similarly shorter for gruesome and aversive stimuli. Women took approximately 0.25 seconds longer (14%) to evaluate the *good* ratings than men, while the *neutral* and *bad* ratings were similar.

### Neuroimaging

The brain activity observed during the neutral condition was subtracted from aversive, erotic, happy, and gruesome conditions, yielding four main comparisons from the study. Overall, the ALC group exhibited lower brain activation values than the NC group for all four contrasts, but significant interactions of group x gender indicated striking differences in these abnormalities. That is, the general observation of lower activation values was evident for ALC_M_, while ALC_W_ exhibited the opposite pattern; the values for each emotion vs. neutral contrast were shifted higher for ALC_W_. **Table 2** identifies regions with significant group x gender interactions for each of the four contrasts. Because the pattern of these significant group x gender interactions was similar for all contrasts, we have chosen to exemplify the two most salient contrasts: erotic vs. neutral **(Figure 3)** and aversive vs. neutral **(Figure 4)**. A summary figure **(Figure 5)** shows the group x gender interactions for all four contrasts.

The contrast of erotic vs. neutral (i.e., erotic minus neutral) is presented in **Figure 3**, which shows that brain activity was greater in most subcortical brain regions for erotic than for neutral images (for ALC_W_, ALC_M_, NC_W_, and NC_M_). The group x gender interaction revealed a significant cluster that encompassed limbic brain regions including the amygdala, thalamus, hippocampus, and parahippocampal cortex, as well as much of the cerebellum. For the medial temporal lobe structures in this cluster, the erotic and neutral pictures elicited less activation difference for ALC_M_ than for NC_M_; this alcoholism-related abnormality was not observed for women.

**Table 2.**
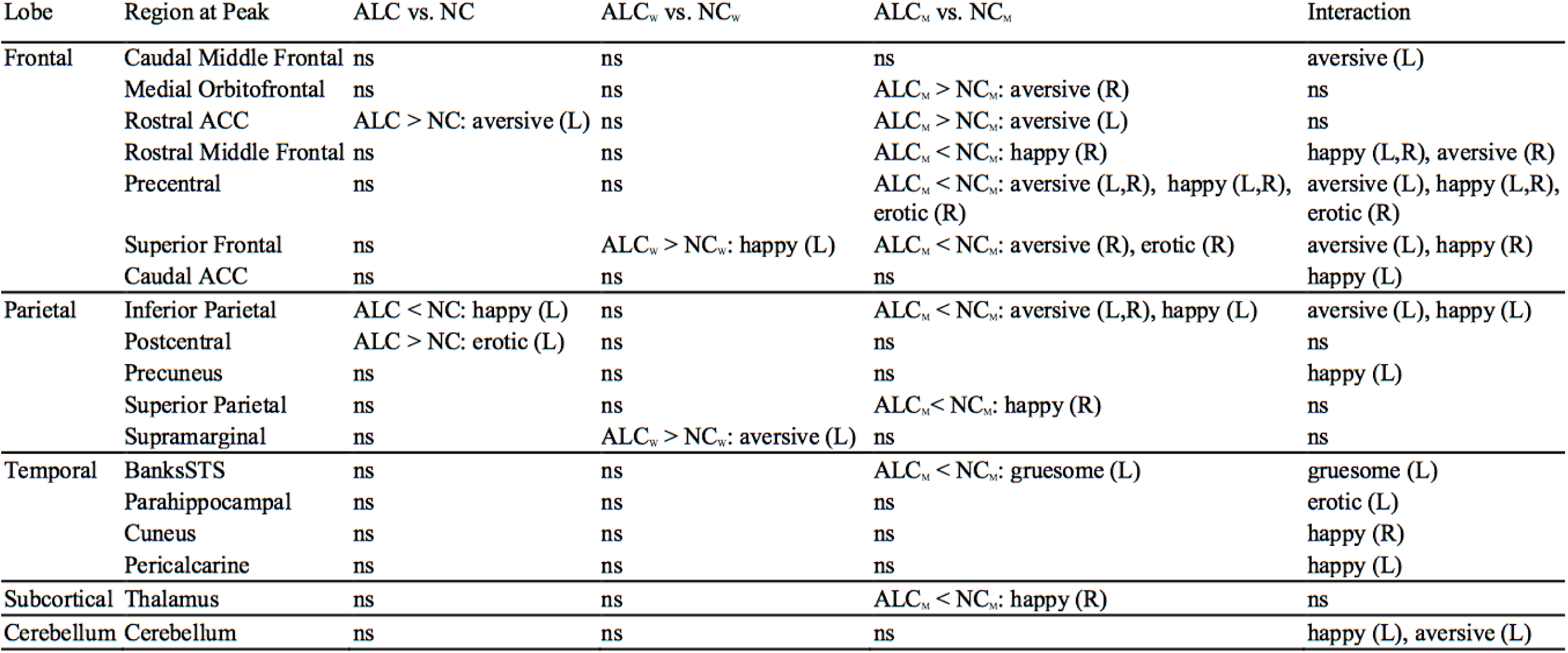
Peak voxel or vertex labels of significant clusters for group contrasts of each emotion vs. neutral condition. Significant dusters (*p* < 0,05 after correction for multiple comparisons) were observed for comparisons between alcoholic and control groups (for the entire sample and for men and women separately), along with group x gender interactions, for each of the four contrasts between each emotion condition compared to the neutral condition. Cortical regions were determined from the peak voxel or vertex. Overall, the table shows that the ALC_M_ had widespread abnormalities in response to emotional stimuli, and that these effects were significantly different than the effects for the ALC_W_. Details are described in the text, **Figure 3, Figure 4**, and **Figure 5**, and in the supplemental tables and figures. Abbreviations: ACC = anterior cingulate cortex; L = left hemisphere; **R** = right hemisphere; ALC_W_ = alcoholic women; ALC_M_ = alcoholic men; NC_W_ = nonalcoholic control women; NC_M_ = nonalcoholic control men; ns = not significant; BanksSTS = banks, superior temporal sulcus.

**Figure 3.**
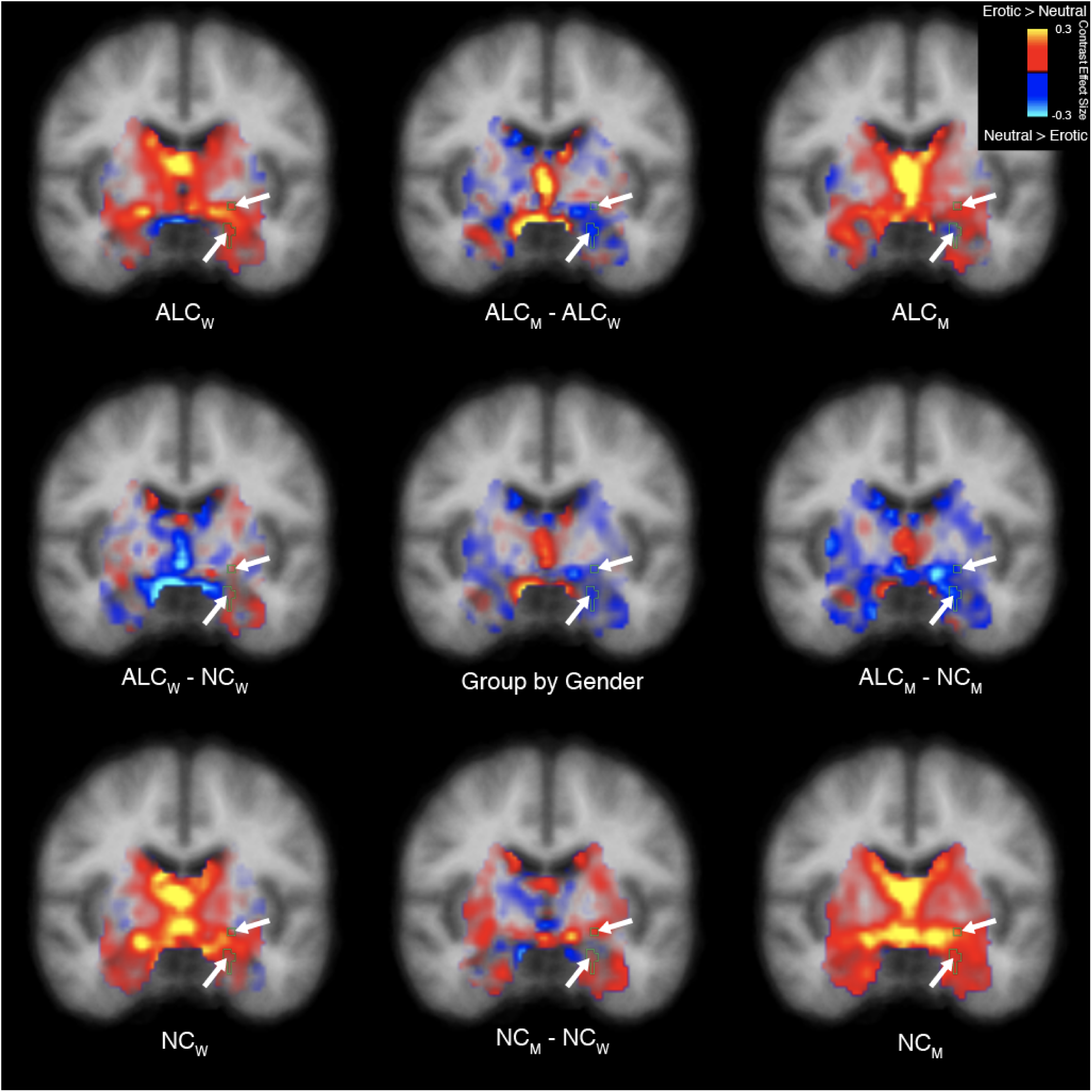
Erotic vs. neutral stimuli elicited abnormal activation of the limbic system and cerebellum in alcoholic men. A significant group x gender interaction in response to erotic vs. neutral stimuli was identified and is displayed as a cyan outline indicated by arrows. The inferior arrow designates the amygdala. Group mean contrast values are displayed in the four brain images located in the corners of the figure, and group comparisons are indicated by minus signs. Contrast values are overlaid on coronal cross sections. Images are displayed in radiological convention with the right hemisphere shown on the left. (Sagittal and axial views are shown in Figure 3-S1 and Figure 3-S2; Figure 3-S3 shows the magnitude of cluster differences.) Abbreviations: ALC_M_ = Alcoholic men; ALC_W_ = Alcoholic women; NC_M_ = Nonalcoholic men; NC_W_ = Nonalcoholic women.

A complex pattern of gender-related alcoholism abnormalities in brain activity were revealed by the contrast of aversive vs. neutral conditions for several significant clusters **(Figure 4)**. For some regions of the brain, activity was higher for aversive than neutral stimuli (“aversive-responding” regions), while for other regions of the brain, activity was higher for neutral than aversive (“neutral-responding” regions). The ALC_M_ - NC_M_ contrast resulted in negative values for both aversive-responding and neutral-responding regions, reflecting the following two situations: For aversive-responding regions, the aversive and neutral stimuli had less activation difference for the ALC_M_ than for the NC_M_, while for neutral-responding regions the aversive and neutral stimuli were more similar for NC_M_ than for the ALC_M_. In four significant clusters, these negative values obtained from ALC_M_ were significantly more negative than those obtained from ALC_W_. As shown in **Figure 4**, three of the clusters were in left prefrontal cortex and one was the inferior parietal gyrus; similar differences were found for the right hemisphere **(Table 2)**. Interestingly, as can be seen in **Table 2**, there also was a significant main effect in two adjoining medial prefrontal regions (medial orbitofrontal and rostral anterior cingulate cortices), wherein alcoholics exhibited higher contrast than controls, and this was more evident in the men than in the women **(Figure 4-S1** and **Figure 4-S3**). For men, this group difference was in the opposite direction as observed for the regions with significant group x gender interactions.

**Figure 4.**
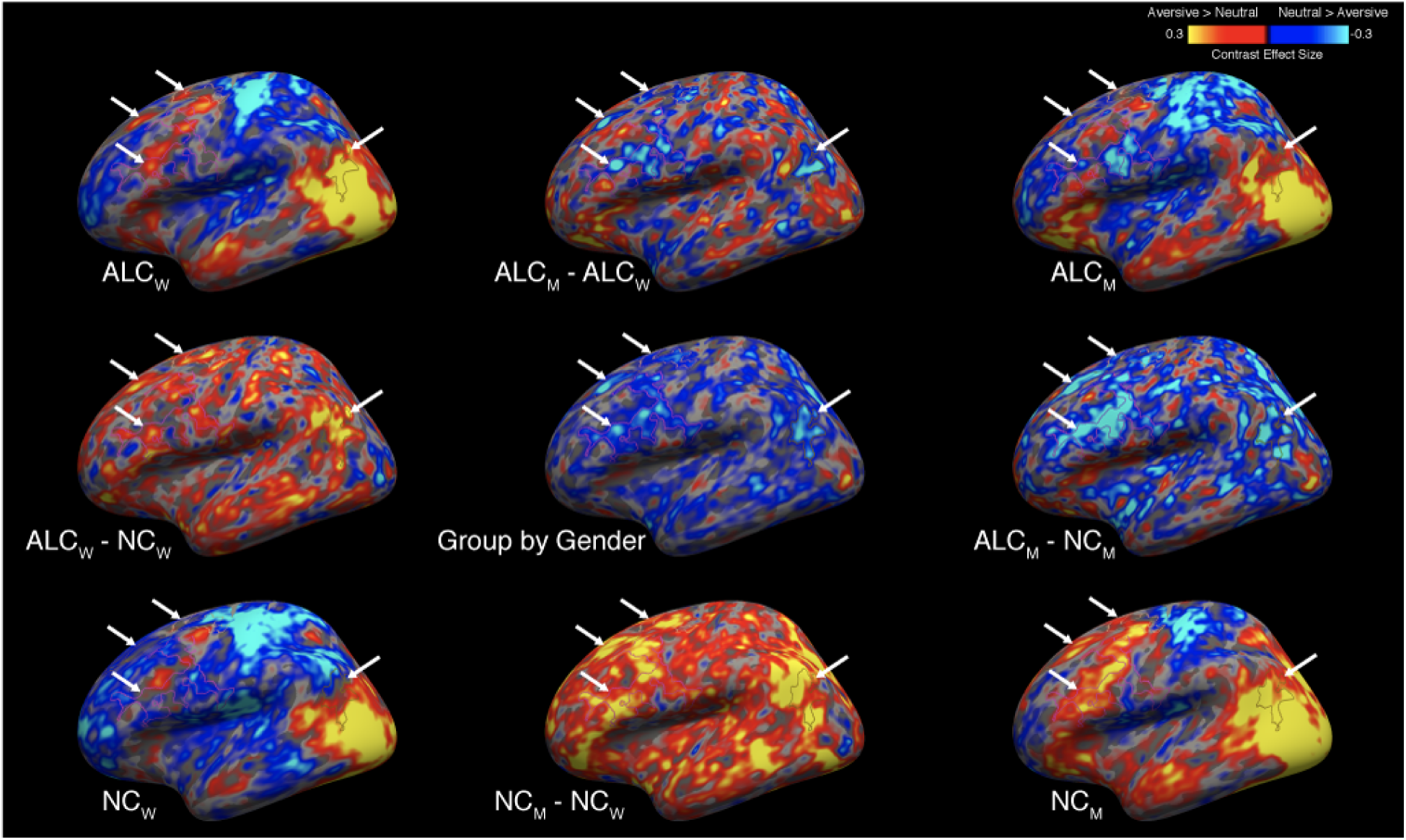
Aversive vs. neutral stimuli elicited more abnormally negative responses in alcoholic men. A significant group x gender interaction revealed several clusters (see Table 2-S2), which are indicated by arrows on the lateral surface of the left hemisphere, overlaid on contrast values between aversive and neutral stimuli. Group mean contrast values (for aversive vs. neutral) are displayed in the four brain images located in the corners of the figure, and group comparisons are indicated by minus signs. (Figure 4-S1 shows the medial surface, while the right hemisphere is shown in Figure 4-S2 for the lateral and Figure 4-S3 for the medial surface; Figure 4-S4 shows the magnitude of cluster differences.). Although not shown here, the activation patterns across the four subgroups for contrasts of other emotional stimuli (i.e., happy, gruesome, and erotic) with neutral stimuli were similar to those shown above, and likewise, the general locations of the activation regions were similar for the four subgroups. Abbreviations: ALC_M_ = Alcoholic men; ALC_W_ = Alcoholic women; NC_M_ = Nonalcoholic men; NC_W_ = Nonalcoholic women.

**Figure 5.**
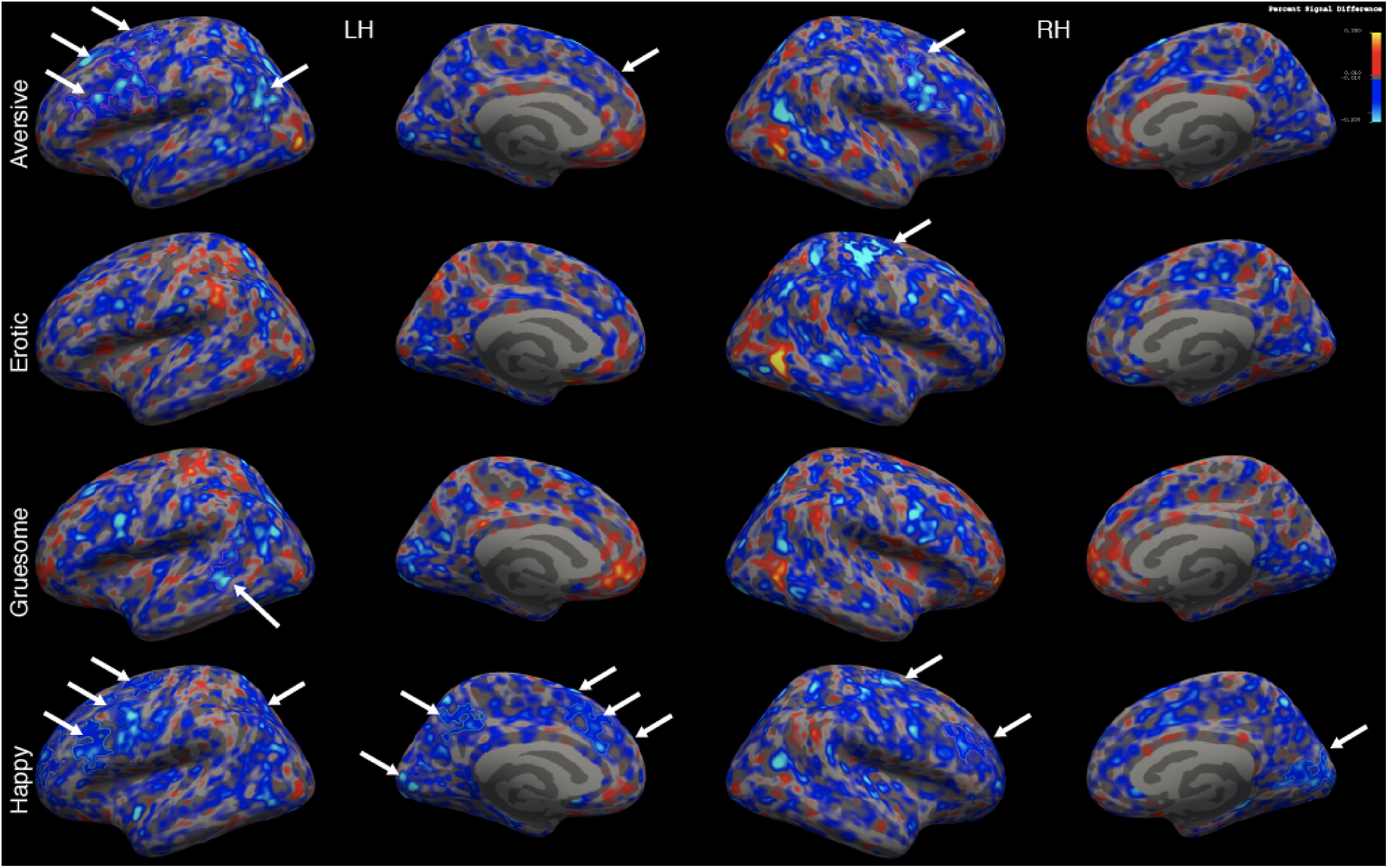
Interaction of group x gender for aversive, erotic, gruesome, and happy stimuli vs. neutral stimuli. Significant clusters are indicated by arrows shown on interaction maps of contrast values for each of the four emotions vs. neutral (similar to the center image in Figure 3 and Figure 4). All four brain surfaces are shown (from left: left lateral, left medial, right lateral, and right medial). Blue regions indicate less activation contrast (emotion vs. neutral) for ALC_M_ relative to NC_M_ vs. ALC_W_ relative to NC_W_. Abbreviations: RH = Right Hemisphere; LH = Left Hemisphere.

In summary, we observed a similar pattern of significant group x gender results **(Figure 5)** for each of the four contrasts (aversive, erotic, gruesome, and happy vs. neutral): ALC_M_ demonstrated less activation for emotional stimuli compared with neutral images, whereas ALC_W_ did not show these decreases and in some contrasts, demonstrated activation increases. For comparison with the observations revealed by the erotic contrast shown in **Figure 3** (which highlights the amygdala) and the aversive contrast shown in **Figure 4** (cortical surface), **Figure 5** shows all four contrasts, including gruesome and happy.

For ALC_W_ compared to NC_W_, significantly more positive brain activation contrasts were seen in superior frontal and supramarginal cortical regions. In ALC_M_ as compared to NC_M_, the contrasts revealed more negative values across widespread areas throughout the brain, including the inferior parietal gyrus, anterior cingulate gyrus, and postcentral gyrus **(Table 2** and **Figure 5)**. Specifically, significant group x gender interactions were observed in the frontal (superior frontal, rostral and caudal middle frontal), parietal (inferior and superior parietal gyri, and precuneus), and occipital (pericalcarine and cuneus) lobes, as well as the caudal anterior cingulate, parahippocampal gyrus, and cerebellum. Happy and aversive contrasts were especially evident throughout widespread regions; the erotic contrast revealed a significant interaction for the parahippocampal gyrus, medial temporal lobe structures, and cerebellum; and the gruesome contrast revealed an interaction for the superior temporal sulcus.

## Discussion

### Alcoholism and Emotional Processing

Deficits in emotional processing contribute to the development and course of alcohol abuse and dependence (Oscar-Berman & Bowirrat, 2005; Oscar-Berman et al., 2014). Research on the relationship between AUD and emotional dysfunction has shown impairments in self-regulation of emotions, as well as deficits in the perception, identification, evaluation, and understanding of emotions of self and others. However, because little is known about the brain responses to emotional stimuli in ALC_W_ as compared to ALC_M_, the present study combined fMRI neuroimaging with a sophisticated experimental design and advanced data analysis methods, to assess the interaction between gender and alcoholism in functional activation of brain regions as participants processed emotional stimuli of varying valences (International Affective Picture System). As indicated in **Table 2**, with the exception of the two ventromedial prefrontal regions, our results showed consistently blunted brain activation contrasts in the ALC group compared to the NC group for men, but this general pattern was not observed for women. Further, a significant interaction between gender and alcoholism indicated that the affective pictures elicited lower activation contrasts in ALC_M_ relative to NC_M_, abnormalities that were significantly lower than those observed between ALC_W_ and NC_W_. In comparison, for ALC_W_, more positive activation contrasts were found in response to emotional stimuli vs. neutral stimuli than found for NC_W_, in regions including the superior frontal and supramarginal cortex.; in ALC_M_, the significant differences were widespread throughout the brain, including the inferior parietal gyrus, anterior cingulate gyrus, and postcentral gyrus. **Table 2** (and supplement) shows the extent and spread of the differences among ALC_M_, ALC_W_, NC_M_, and NC_W_.

### Gender and Alcoholism Interaction in Emotional Processing Regions in the Brain

Emotional processing involves engaging multiple brain regions (Oscar-Berman & Bowirrat, 2005). *In vivo* neuroimaging studies as well as *post-mortem* pathological studies have shown that cortical loss in the frontal lobes is the most common damage observed both in association with AUD (Oscar-Berman & Marinkovic, 2003) and in individuals having emotional disorders unrelated to AUD (Bechara, Damasio, & Damasio, 2000; Young et al., 2010). In our study, the regions showing significant interactions between alcoholism and gender were mainly in the frontal lobes: precentral, rostral and caudal middle frontal cortex, superior frontal cortex, and the caudal anterior cingulate cortex for both happy and aversive stimuli. Previous fMRI studies have suggested that rostral middle frontal cortex may be involved in the implicit or uninstructed generation and perpetuation of emotional states (Waugh, Hamilton, & Gotlib, 2010; Waugh, Lemus, & Gotlib, 2014). Given that in our study, ALC_M_ showed lower activation in the rostral middle frontal cortex in response to happy stimulation vs. neutral stimulation, our finding may reflect deficits in ALC_M_ in maintaining positive emotion in response to happy stimuli. Aldhafeeri and colleagues (Aldhafeeri, Mackenzie, Kay, Alghamdi, & Sluming, 2012) reported significant activation in the prefrontal cortex, anterior and posterior cingulate gyri, and temporal lobes in healthy volunteers, in response to pleasant pictures vs. neutral pictures.

One of the other frontal brain regions that showed a significant gender x alcoholism interaction was the caudal anterior cingulate cortex, a region thought to be involved in appraisal and expression of negative emotion (Etkin, Egner, & Kalisch, 2011). However, for the regions anterior to the caudal anterior cingulate, we found a different pattern of group differences. The ALC group had greater contrast values than the NC group (especially evident for ALC_M_) in the subcallosal regions of medial orbitofrontal cortex and rostral anterior cingulate cortex. For the ALC_M_, the difference was in the opposite direction to that observed for other widespread regions, where group x gender interactions had been evident. Because these ventromedial prefrontal regions are involved in the integration of cognitive and affective responses to external events (Bush, Luu, & Posner, 2000; Margulies et al., 2007), the increased responsivity in the ALC group might indicate compensatory involvement in evaluating the emotional pictures (Oscar-Berman & Marinković, 2007). Additionally, consideration of the possibility that these ventromedial prefrontal regions are associated with hyperactivity to IAPS stimulus contrasts in other psychopathological conditions (Hägele et al., 2016).

Additionally, significant interactions between gender and alcoholism were observed in cortical regions involved mainly in visual processing, including the cuneus and precalcarine regions, in response to happy stimuli **(Figure 5)**. These significant interactions reflected higher BOLD values to affective pictures compared to neutral pictures, more so in NC_M_ than ALC_M_, whereas the effect was reduced or reversed for the two groups of women. In NC participants, we confirmed the greater activation in visual cortex while viewing emotional vs. neutral pictures that has been reported in prior studies, with some suggesting stronger responses by men to pleasant pictures and stronger responses by women to unpleasant pictures (Sabatinelli, Flaisch, Bradley, Fitzsimmons, & Lang, 2004).

Inferior parietal cortex was another region that showed a significant interaction between gender and alcoholism, driven mainly by the blunted activation in the ALC_M_ compared to the NC_M_ men. Inferior parietal gyrus is involved in the perception of emotions in facial stimuli (Sarkheil, Goebel, Schneider, & Mathiak, 2013). Except for neutral pictures, most of the other pictures had a human face in them, and therefore, the interaction and lower activation in ALC_M_ may represent an impairment in processing emotional facial expressions. In fact, previous studies have shown that abstinent ALC_M_ showed less activation in temporal limbic areas, when viewing positive or negative emotional faces compared to controls (Marinkovic et al., 2009).

There also were significant interactions between gender and alcoholism in limbic and subcortical structures: In ALC_M_, brain activity for erotic and neutral pictures were relatively similar leading to decreased differential activation, while NC_M_ had stronger activity for erotic than neutral pictures, for parahippocampal cortex, hippocampus, amygdala, other limbic structures, and the cerebellum. This alcoholism-related abnormality was not observed for women: ALC_W_ had a slightly larger (although not significant) positive contrast between erotic and neutral pictures compared to NC_W_. Limbic structures are involved in emotional processing and decision making, and the muted responsivity observed for ALC_M_ could reflect deficits in these functions.

### Potential Mechanisms

Excessive alcohol consumption can facilitate widespread physical deterioration, including neurodegeneration (Cargiulo, 2007). However, due to the heterogeneity of AUD etiology, the risks and extent of impairments as well as treatment outcomes can vary widely (Soyka & Schmidt, 2009). It is likely that environmental influences, psychological vulnerabilities, and individual differences such as genetic variation, drinking history, and body composition account for much of this variance, making certain subgroups, such as women, more vulnerable to the effects of alcohol (Brady & Sonne, 1999; Mareckova et al., 2016; Nixon et al., 2014; Oscar-Berman & Marinkovic, 2007). As impairments in emotional response and corresponding brain dysfunction may be central to the etiology of AUD, delineating the relationship between gender and emotional reactivity in relation to alcoholism increases our understanding of its etiology and affords the opportunity to create novel interventions tailored to the presentation of individual patients.

The present study is consistent with the view that women and men may develop AUD for different reasons (Erol & Karpyak, 2015), with women seeking to dampen aversive emotional reactivity and men desiring to feel more enlivened and social (Mosher Ruiz et al., 2017). Moreover, psychiatric conditions such as depression in conjunction with AUD have a higher prevalence in women than in men (Khan et al., 2013), whereas antisocial personality has been observed in conjunction with AUD more often in men (Compton, Conway, Stinson, Colliver, & Grant, 2005; Merikangas et al., 1996; Oscar-Berman et al., 2009).

A diagnosis of AUD often is associated with comorbid psychiatric illnesses characterized by dysregulated affect. Given the relationship between emotional dysregulation and other psychiatric disorders (e.g., Generalized Anxiety Disorder, Major Depressive Disorder, and Antisocial Personality Disorder) (Richard J. Davidson, Pizzagalli, Nitschke, & Putnam, 2002; Regier et al., 1990), the present findings also suggest that the patterns of neural reactivity observed in our study may contribute to the onset of multiple diagnostically distinct syndromes. Research that could further elucidate shared aspects of these disorders are particularly relevant for clinical care, as diagnostic comorbidity in alcoholics is associated with an increased risk for suicide (Driessen et al., 1998; Regier et al., 1990). Finally, while there are clear gender differences in drinking preferences among college students (Labrie, Cail, Hummer, Lac, & Neighbors, 2009), and women on average display increased emotional reactivity relative to men (Proverbio et al., 2009), the present findings highlight these effects in long-term abstinent alcoholics.

The results of this study are to be considered in the context of several limitations. First, our results are based upon cross-sectional data, and as such, it is impossible to determine if chronic alcohol usage caused, or resulted from, the observed dysregulated emotional reactivity, or perhaps a combination of both. Second, we had limited information about the potentially confounding variable of smoking status, and therefore, it was not included in the analyses. Smoking abstinence has been associated with increased emotional reactivity in response to unpleasant stimuli (Versace et al., 2012) and interactions with alcoholism (Durazzo et al., 2013; Luhar, Sawyer, Gravitz, Ruiz, & Oscar-Berman, 2013), and therefore, may have influenced the results of the present study. Third, while there were peak regions of activation differences, these were observed against a background of broad regions identified that were different between each of the emotional conditions and the neutral condition, and the significant group x gender interactions reflected these broad differences in brain activity. We chose not to artificially suppress the display of these widespread effects in our figures by restricting the thresholds. Despite these considerations, the present findings highlight the need for continued research on the overlap between gender differences in processing of emotional stimuli and the development of pathological alcohol consumption.

## Conclusions

While blunted emotional reactivity had been observed previously in alcoholics, earlier studies had focused either exclusively on men or had collapsed data across genders (Gilman, Davis, & Hommer, 2010; Marinkovic et al., 2009; Salloum et al., 2007). Therefore, the present study provides additional insights into emotional processing in alcoholism by examining the influence of gender on brain activation. In our previous studies (Rivas-Grajales et al., 2018; Sawyer et al., 2018, 2017, 2016; Seitz et al., 2016), we had reported gender differences in morphometry of cerebral and cerebellar subregions, and white matter integrity, in association with alcoholism history in men and women. In the current study, we reported functional abnormalities in cortical, subcortical, and cerebellar regions involved in emotional processing that were different in alcoholic men and women. Significant interactions between alcoholism and gender in several cortical regions in response to emotional stimuli were observed for the aversive and happy stimuli, as well as large differences between ALC_M_ and NC_M_. Areas within the frontal lobes were among the brain regions evidencing the most profound alcoholism-related gender differences.

The brain activity contrasts related to affective vs. neutral stimuli were dampened in alcoholic men in the current study, similarly to prior research showing that ALC_M_ had blunted limbic activation to emotionally expressive faces (Marinkovic et al., 2009). Women are traditionally believed to be more emotionally reactive than men (Merikangas et al., 1996), and in the current study, whereas ALC_M_ had decreased fMRI emotional responsivity, ALC_W_ had similar or greater brain activity in response to emotional stimuli than NC_W_, leading to significant group by gender interaction effects. Our findings support the view that alcohol can be abused in an effort to restore emotional homeostasis either by increasing or decreasing emotional arousal. Future research should prospectively examine gender differences in emotional reactivity and subsequent drinking behavior, to determine the contributions of gender differences that precede AUD, as compared to gender differences that develop as a result of chronic alcoholism.

## Acknowledgements

This research was supported by US Department of Veterans Affairs Clinical Science Research and Development (I01CX000326); National Institute on Alcohol Abuse and Alcoholism (NIAAA) of the National Institutes of Health US Department of Health and Human Services (R01AA07112, R01AA016624, K05AA00219, and K01AA13402); Athinoula A. Martinos Center for Biomedical Imaging Shared Instrumentation Grants (1S10RR023401, 1S10RR019307, and 1S10RR023043); Alcoholic Beverage Medical Research Foundation; Mental Illness and Neuroscience Discovery (MIND) Institute; NIH National Center for Research (P41RR14075); and Boston University Clinical and Translational Sciences Institute (BU CTSI; 1UL1TR001430). We gratefully thank Elinor Artsy, Sheeva Azma, Julie Howard, Sharon Jaffin, Chris Markiewicz, Diane Merritt, Alan Poey, Daniel Salz, Yulia Spantchak, Maria Valmas, and Robert Zondervan for help with recruitment assistance, materials, data collection, consultation, and analysis. The content is solely the responsibility of the authors and does not necessarily represent the official views of the National Institutes of Health, the U.S. Department of Veterans Affairs, or the United States Government.

## Supplemental tables and figures

**Table 1S-1.**
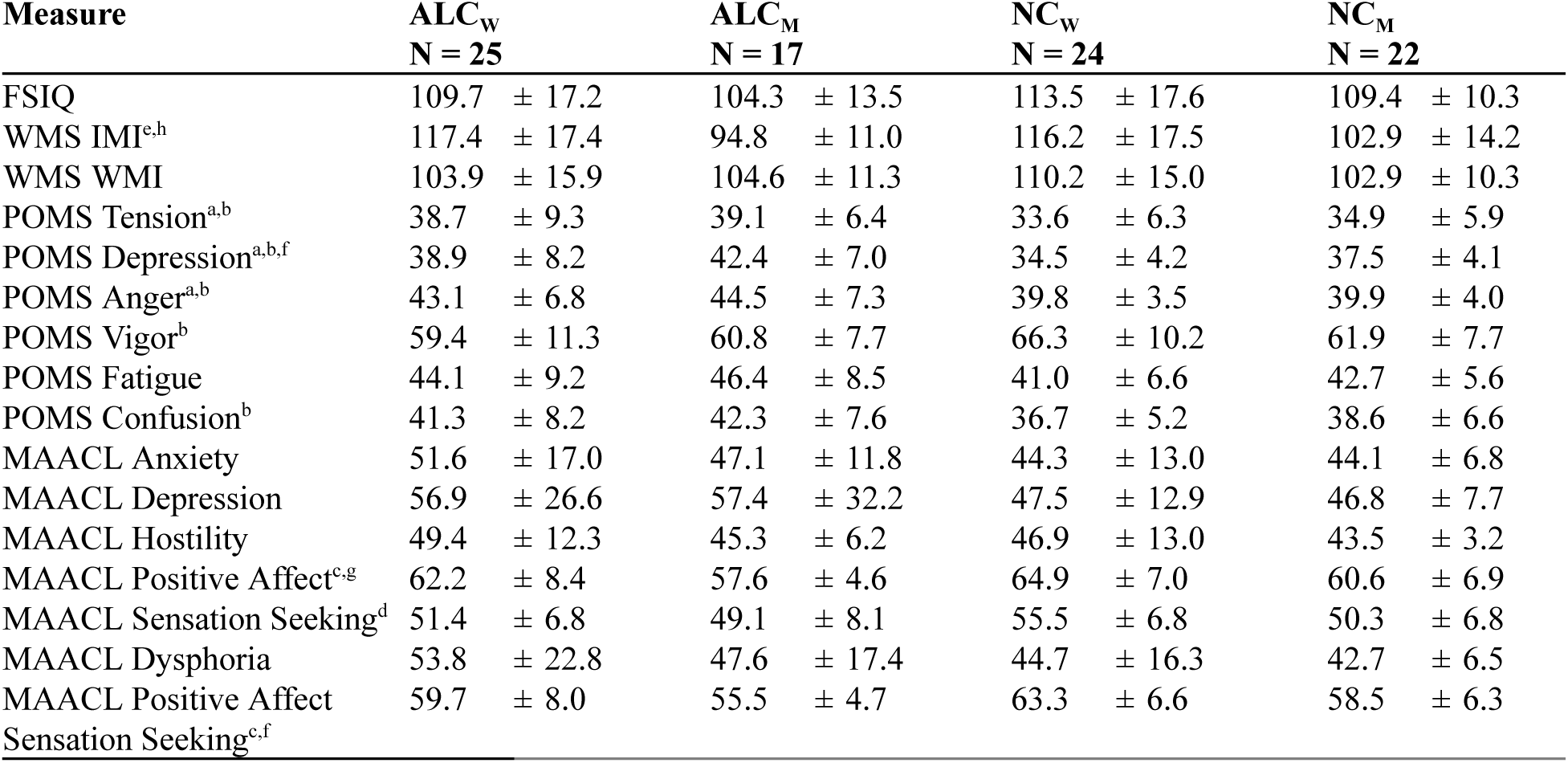
Neuropsychological and affect scores for alcoholic men and women. Abbreviations: ALC_W_ = alcoholic women; ALC_M_ = alcoholic men; NC_W_ = nonalcoholic control women; NC_M_ = nonalcoholic control men; WMS DMI = Wechsler Memory Scale, 3rd ed. Delayed (General) Memory Index; POMS = Profile of Mood States Manual (McNair, 1971); MAACL = Multiple Affective Adjective Checklist (Zuckerman & Lubin, 1985). Significant differences: ^a^(ALC_M_ > NC_M_, *p*<0.05): ^b^(ALC_W_ > NC_W_, *p*<0.05); ^c^(ALC_W_ > ALC_M_, *p*<0.05): ^d^(ALC_W_ < ALC_M_, *p*<0.05); ^e^(ALC_M_ < ALC_W_, *p*<0.001); ^f^(NC_M_ > NC_W_, *p*<0.05); ^g^(NC_M_ < NC_W_, *p*<0.05); ^h^(NC_M_ < NC_W_, *p*<0.01).

**Figure 2-S1.**
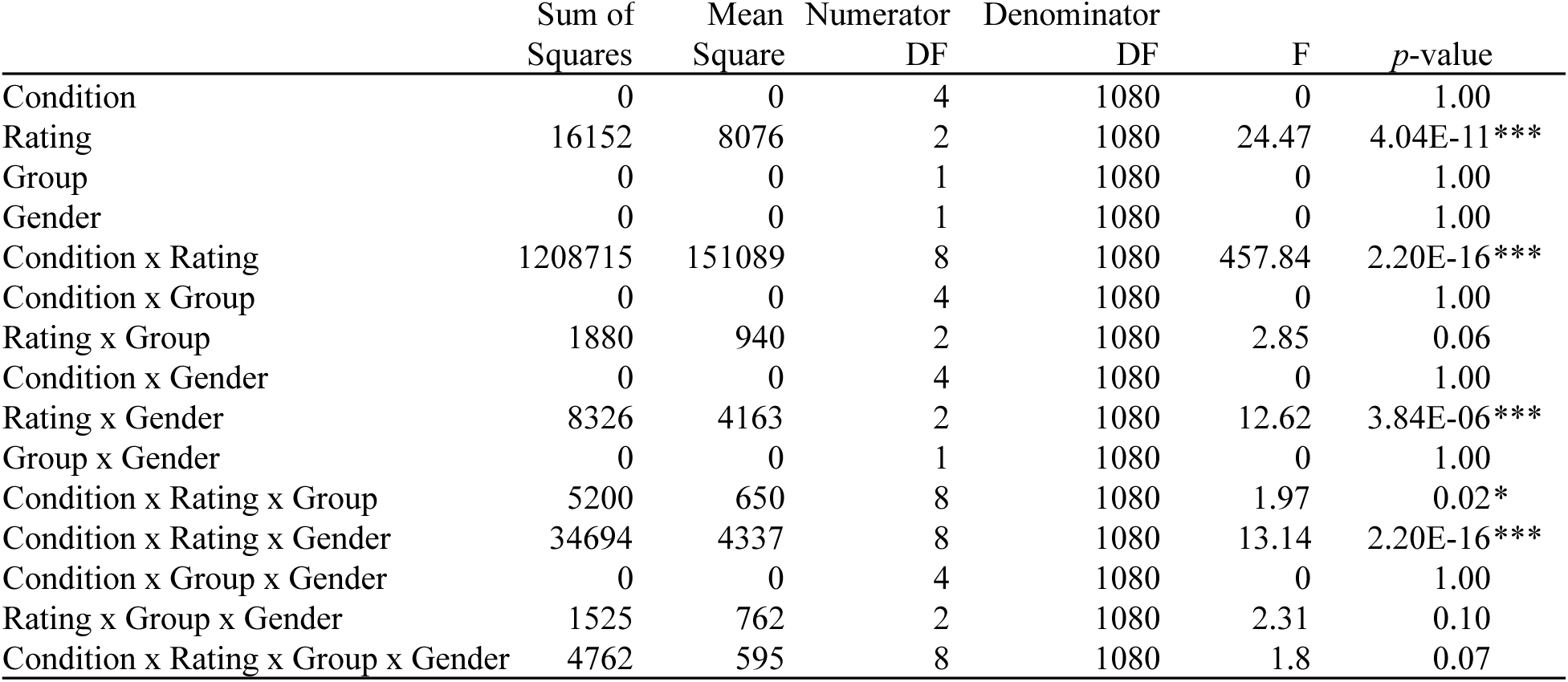
Analysis of variance for percentage of pictures rated. Abbreviations: DF = degrees of freedom. Significance codes: \*\*\**p*<0.001; \**p*<0.05

**Figure 2-S2.**
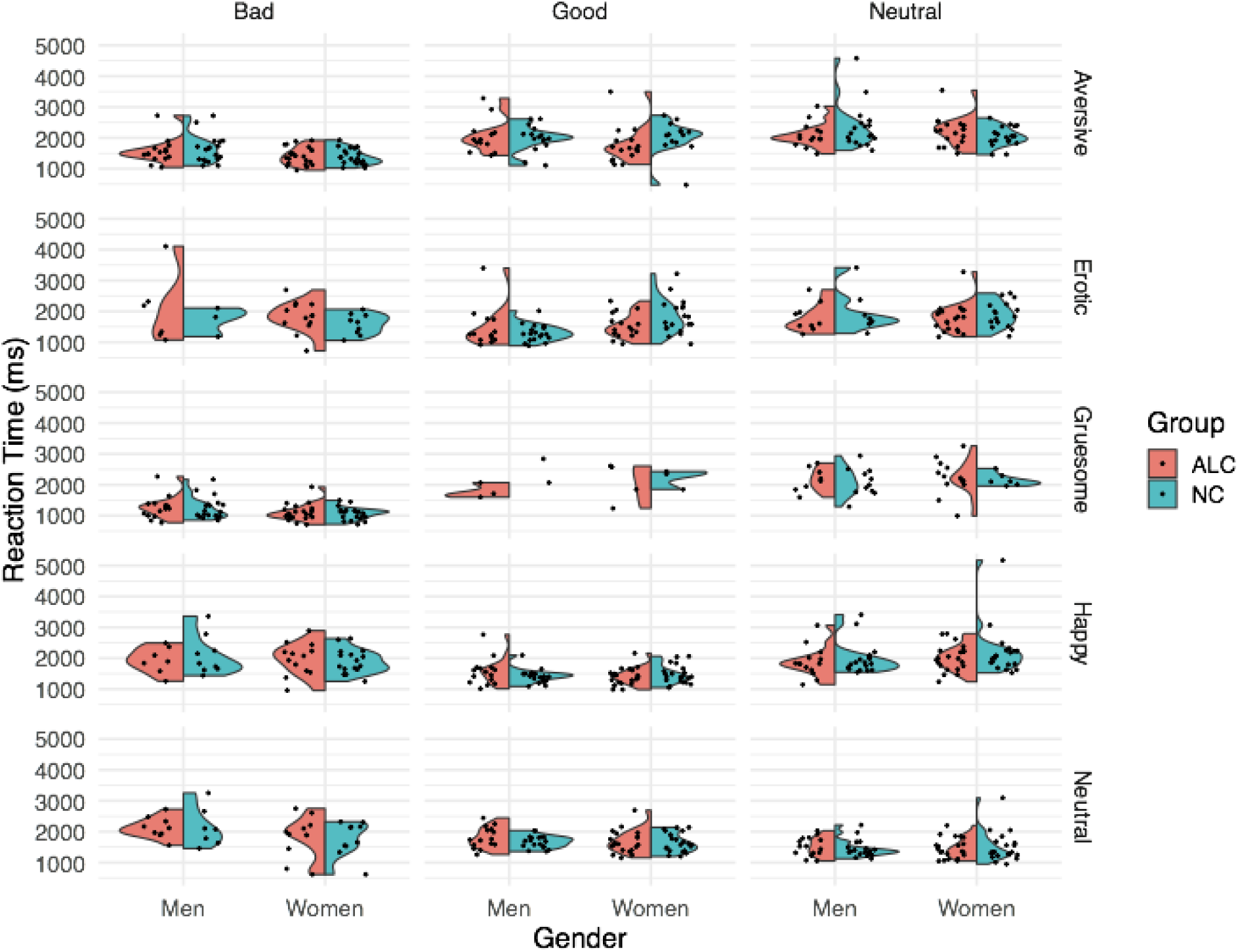
Reaction times of behavioral ratings by condition, rating, group, and gender. The split violin plot represents the significant rating x gender interaction for reaction times of the pictures *p*<0.05 (Figure 2-S3). The difference in the *good* rating of the erotic pictures by alcoholics vs. nonalcoholic controls. The ratings for other conditions were qualitatively similar for alcoholics and nonalcoholic control subjects. Figure 2 shows the reaction times. ALC = alcoholic participants; NC = nonalcoholic control participants.

**Figure 2-S3.**
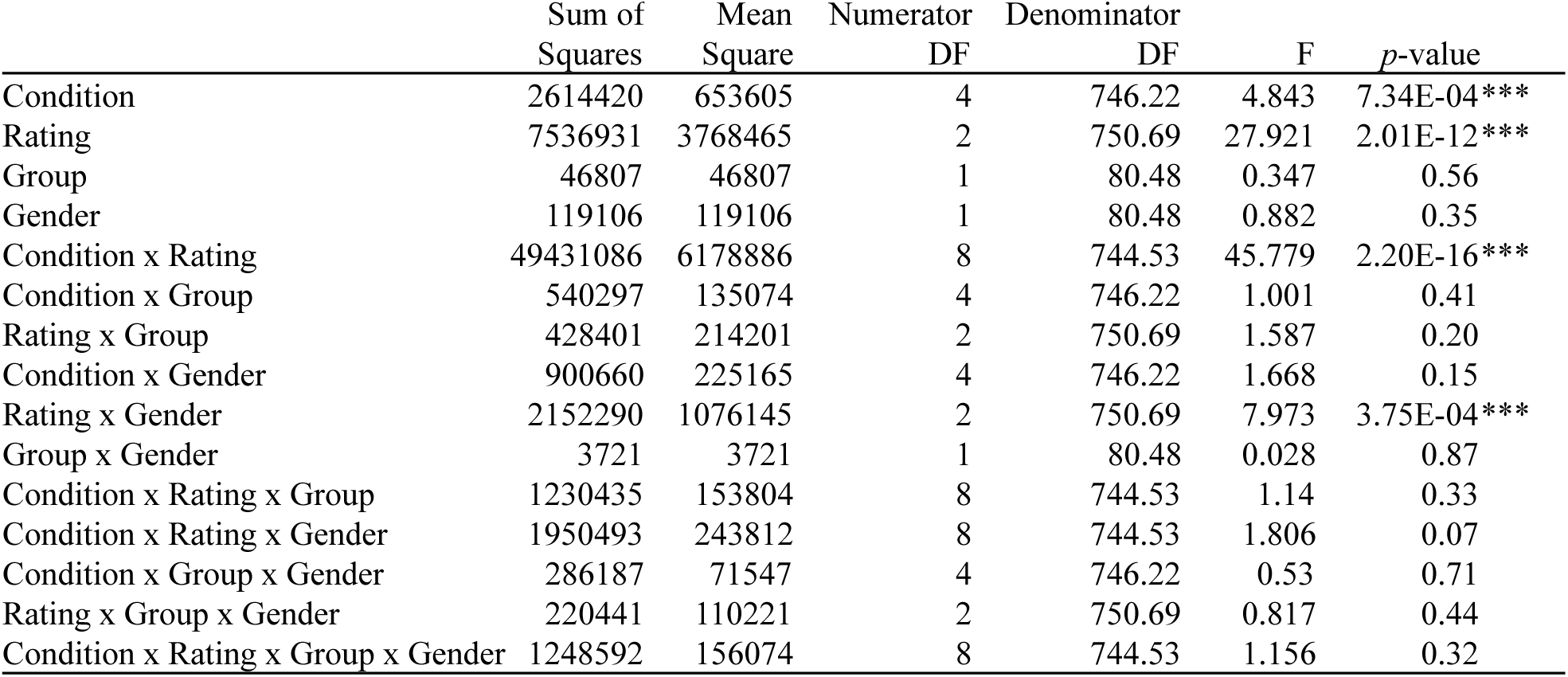
Analysis of variance for reaction times of pictures rated. Abbreviations: DF = degrees of freedom. Significance codes: \*\*\**p*<0.001

**Figure 3-S1.**
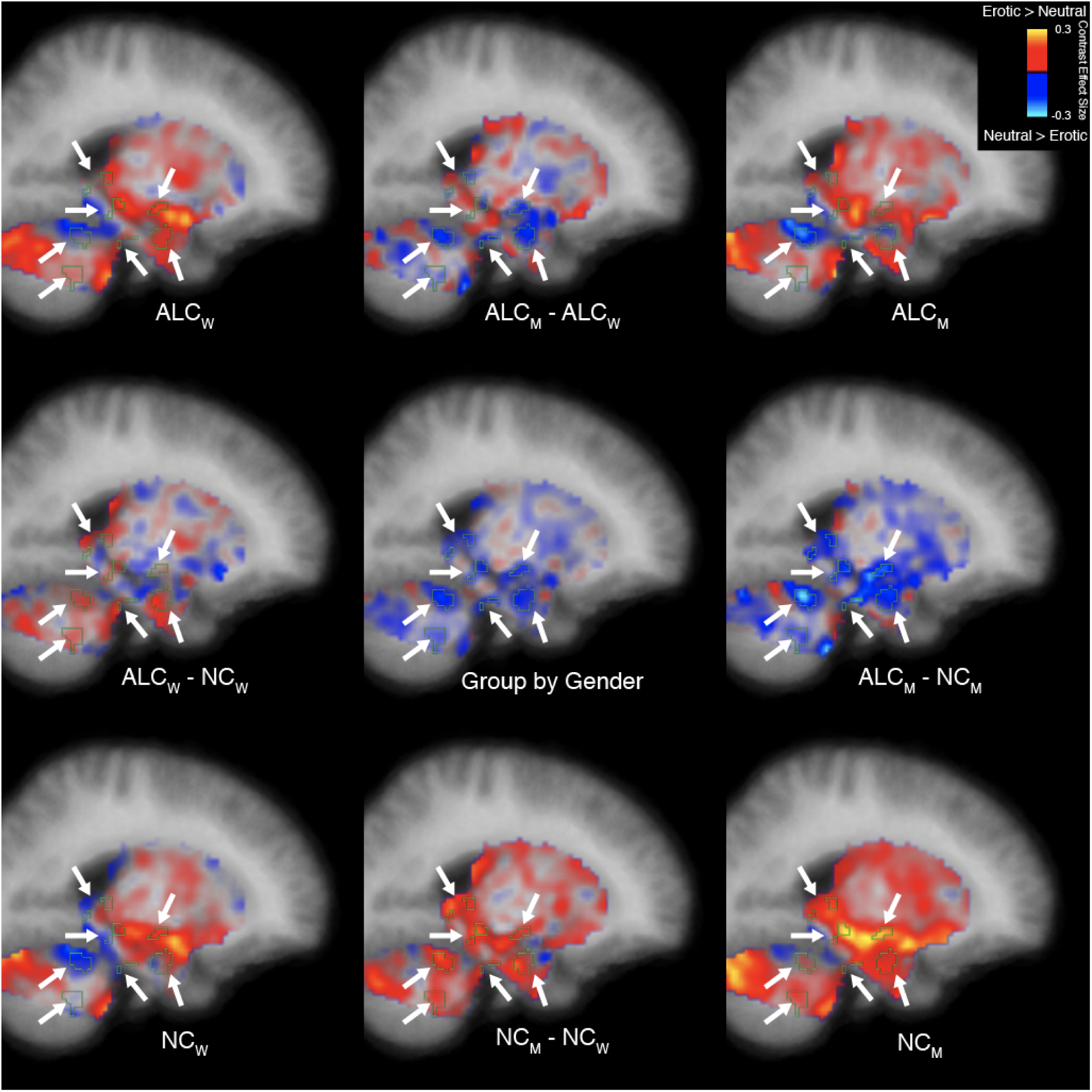
Erotic vs. neutral stimuli elicited abnormal activation of the limbic system and cerebellum in alcoholic men (sagittal view). A group x gender interaction in response to erotic vs. neutral stimulation was identified and is displayed as a thin cyan outline. Group mean values are displayed in the four brain images located in the corners of the figure, and group comparisons are indicated by minus signs. Contrast effect sizes are overlaid on sagittal cross sections. (Coronal and axial views are shown in Figure 3 and Figure 3-S2; Figure 3-S3 shows the magnitude of cluster differences.) Abbreviations: ALC_M_ = alcoholic men; ALC_W_ = alcoholic women; NC_M_ = nonalcoholic men; NC_W_ = nonalcoholic women.

**Figure 3-S2.**
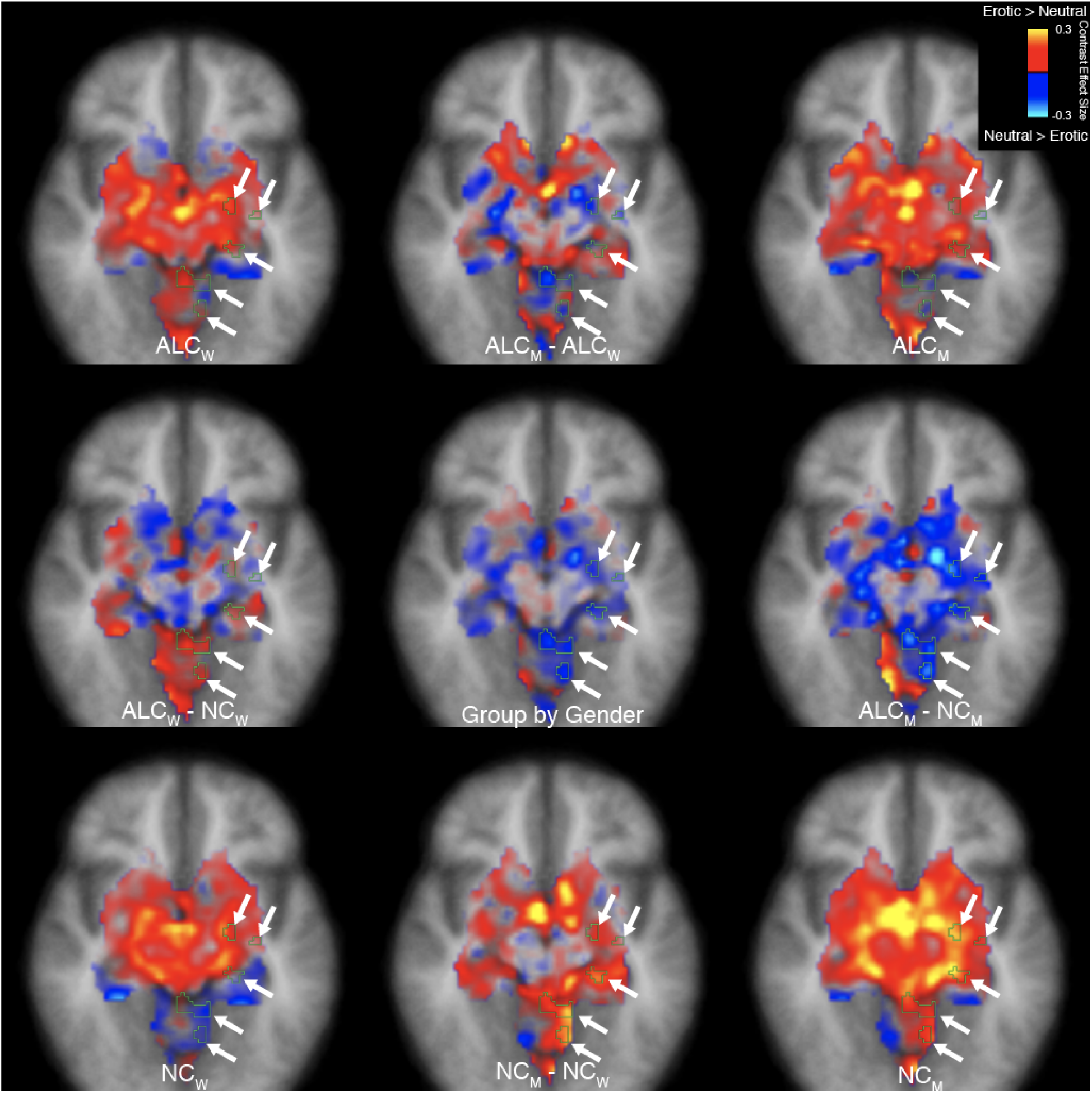
Erotic vs. neutral stimuli elicited abnormal activation of the limbic system and cerebellum in alcoholic men (axial view). A group x gender interaction in response to erotic vs. neutral stimulation was identified and is displayed as a thin cyan outline. Group mean values are displayed in the four brain images located in the corners of the figure, and group comparisons are indicated by minus signs. Contrast effect sizes are overlaid on axial cross sections. Images are displayed in radiological convention with the right hemisphere shown on the left. (Coronal and sagittal views are shown in Figure 3 and Figure 3-S1; Figure 3-S3 shows the magnitude of cluster differences.) Abbreviations: ALC_M_ = alcoholic men; ALC_W_ = alcoholic women; NC_M_ = nonalcoholic men; NC_W_ = nonalcoholic women.

**Figure 3-S3.**
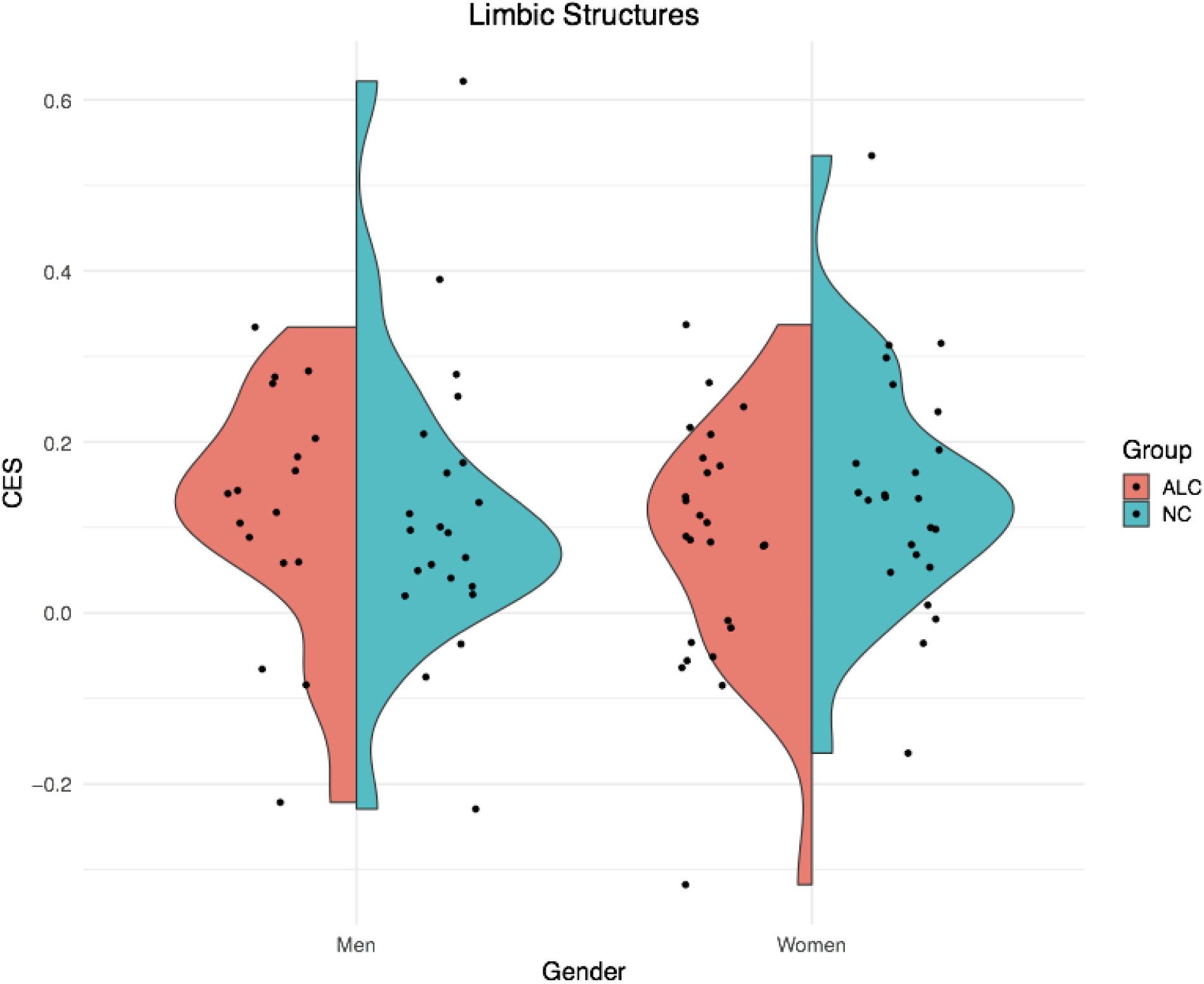
Contrast values observed in the cluster for erotic vs. neutral conditions. The split violin plot represents the Contrast Effect Size (CES; equivalent to “CON” in SPM or “COPE” in FSL) for the cluster in which a significant group x gender interaction was identified for the erotic vs. neutral contrast (*p*<0.05 after correction for multiple comparisons). Positive values indicate erotic > neutral, while negative values indicate erotic < neutral. Each point represents a single participant’s average CES for vertices within the cluster. This figure is meant to convey the variability in CES across participants that is not visible in Figure 3. Nonalcoholic control men had greater activation to erotic stimuli than neutral stimuli, and the contrast was more positive than was observed for alcoholic men. The pattern was reversed for women: alcoholic women had lower contrast values than nonalcoholic women. Abbreviations: ALC = alcoholic participants; NC = nonalcoholic control participants.

**Figure 4-S1.**
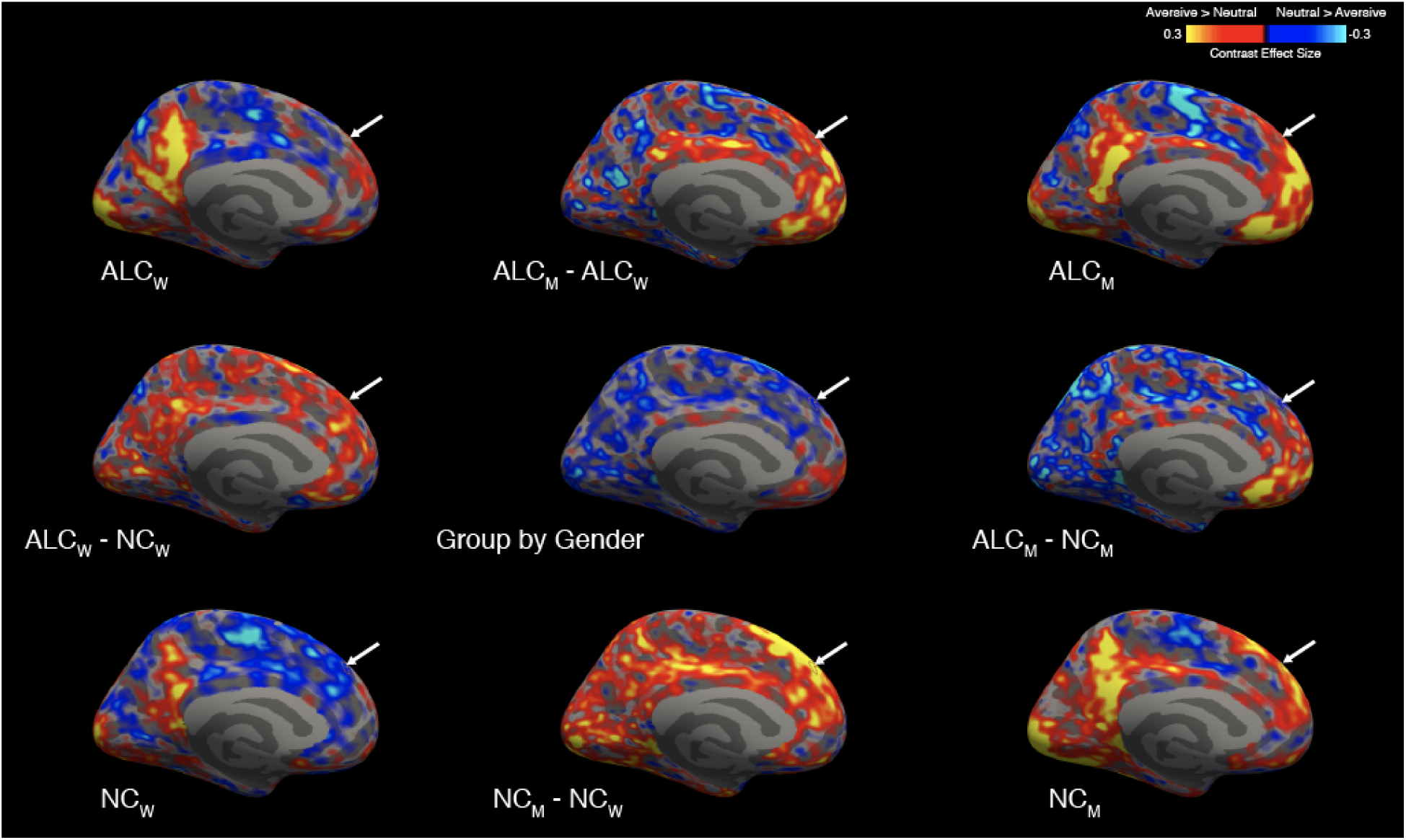
Aversive vs. neutral stimuli elicited more abnormally negative responses in alcoholic men (left medial surface). A group x gender interaction revealed several clusters (see Table 2-S2), which are displayed as thin outlines and indicated by arrows on the medial surface of the left hemisphere, overlaid on contrast values between aversive and neutral stimuli. Group mean values (for aversive vs. neutral) are displayed in the four brain images located in the corners of the figure, and group comparisons are indicated by minus signs. (Figure 4 shows the lateral surface, while the right hemisphere is shown in Figure 4-S2 for the lateral and Figure 4-S3 for the medial surface; Figure 4-S4 shows the magnitude of cluster differences.). Although not shown here, the activation patterns across the four subgroups for contrasts of other emotional stimuli (i.e., happy, gruesome, and erotic) with neutral stimuli were similar to those shown above, and likewise, the general locations of the activation regions were similar for the four subgroups. Abbreviations: ALC_M_ = alcoholic men; ALC_W_ = alcoholic women; NC_M_ = nonalcoholic men; NC_W_ = nonalcoholic women.

**Figure 4-S2.**
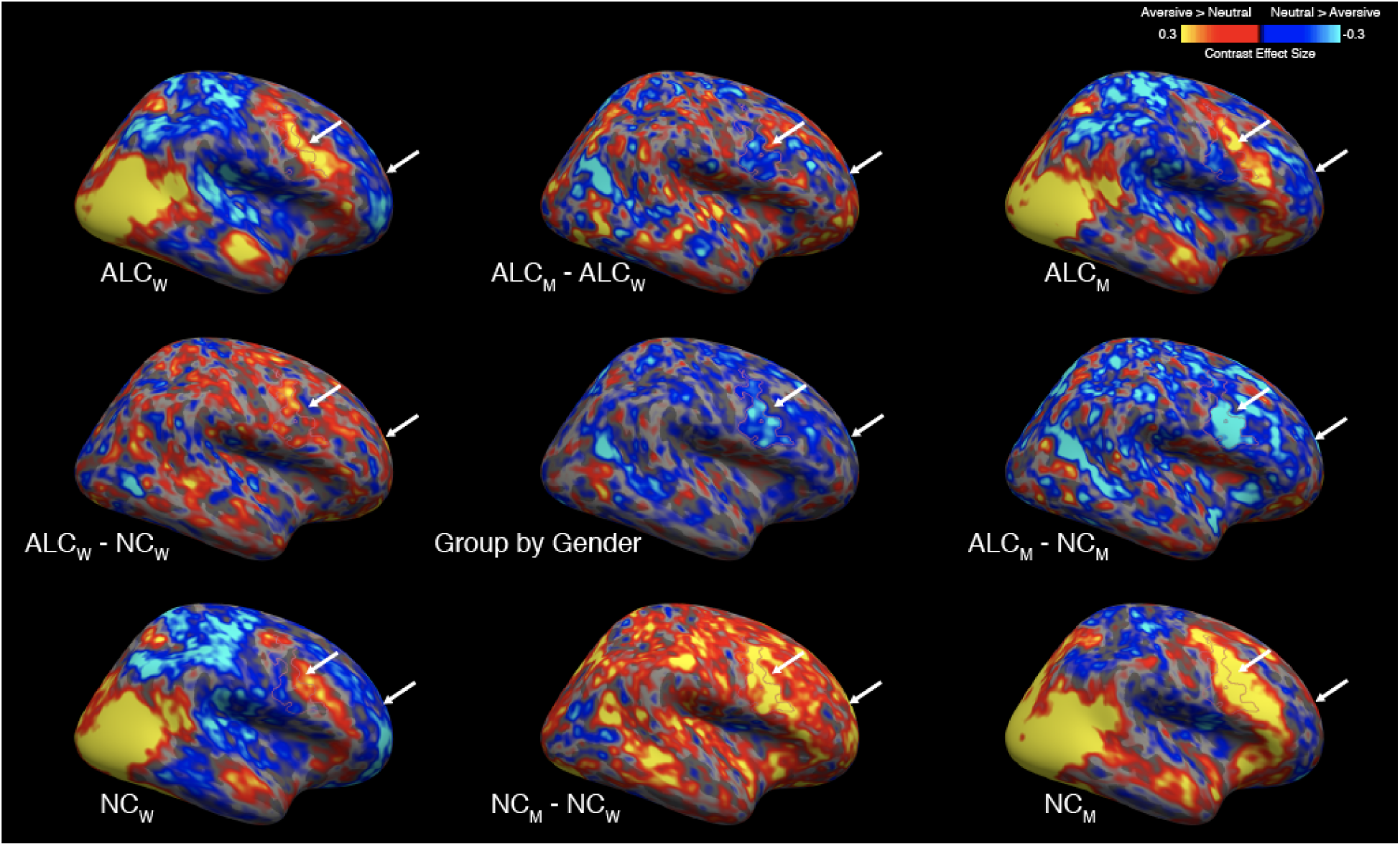
Aversive vs. neutral stimuli elicited more abnormally negative responses in alcoholic men (right lateral surface). A group x gender interaction revealed several clusters (see Table 2-S2), which are displayed as thin outlines and indicated by arrows on the lateral surface of the right hemisphere, overlaid on contrast values between aversive and neutral stimuli. Group mean values (for aversive vs. neutral) are displayed in the brain images located in the four corners of the figure, and group comparisons are indicated by minus signs. (Figure 4 and Figure 4-S1 show the lateral and medial surfaces of the left hemisphere, while the medial surface of the right hemisphere is shown in Figure 4-S3; Figure 4-S4 shows the magnitude of cluster differences.). Although not shown here, the activation patterns across the four subgroups for contrasts of other emotional stimuli (i.e., happy, gruesome, and erotic) with neutral stimuli were similar to those shown above, and likewise, the general locations of the activation regions were similar for the four subgroups. Abbreviations: ALC_M_ = alcoholic men; ALC_W_ = alcoholic women; NC_M_ = nonalcoholic men; NC_W_ = nonalcoholic women.

**Figure 4-S3.**
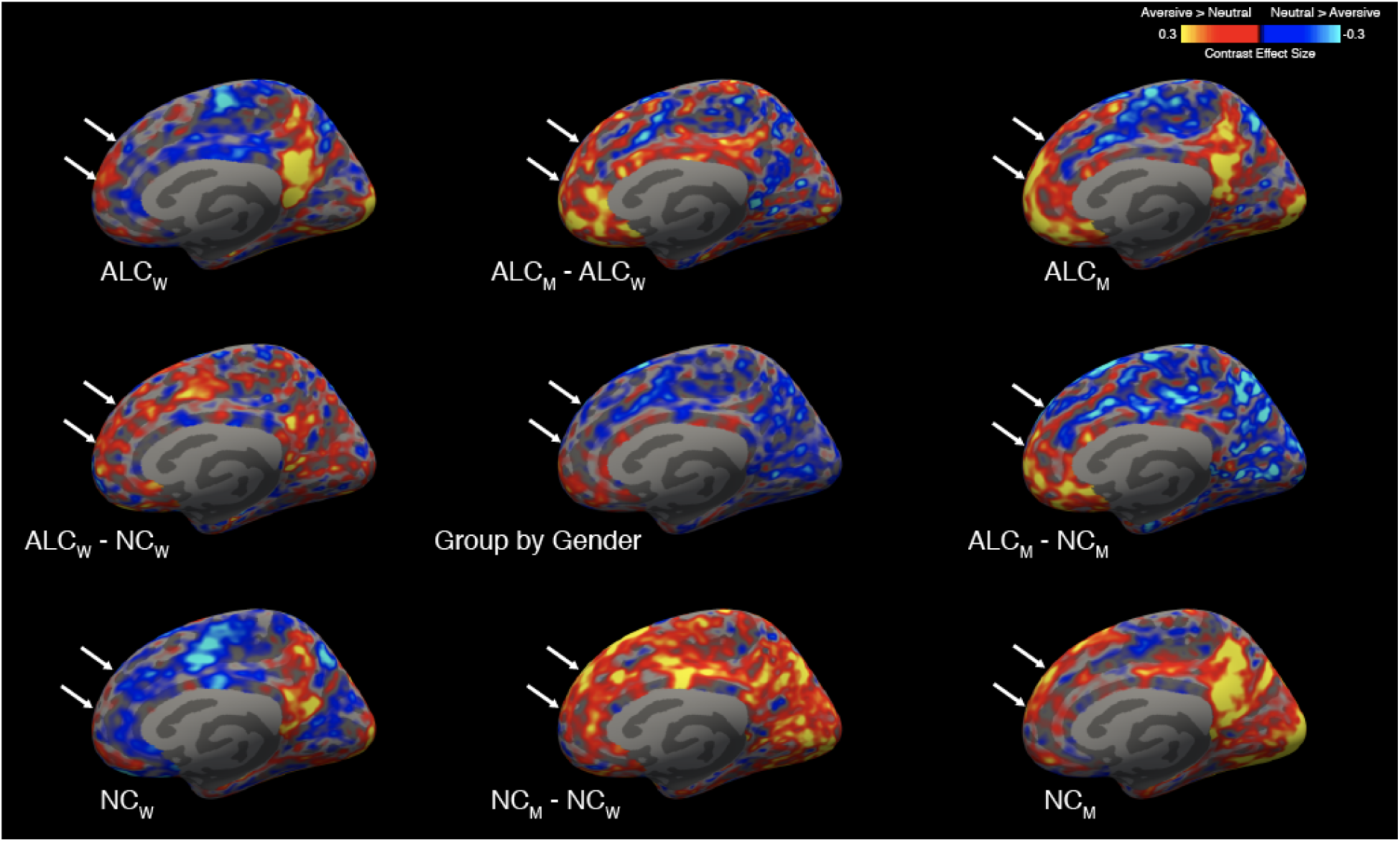
Aversive vs. neutral stimuli elicited more abnormally negative responses in alcoholic men (right medial surface). A group x gender interaction revealed several clusters (see Table 2-S2), which are displayed as thin outlines and indicated by arrows on the medial surface of the right hemisphere, overlaid on contrast values between aversive and neutral stimuli. Group mean values (for aversive vs. neutral) are displayed in the brain images located in the four corners of the figure, and group comparisons are indicated by minus signs. (Figure 4 and Figure 4-S1 show the lateral and medial surfaces of the left hemisphere, while the lateral surface of the right hemisphere is shown in Figure 4-S2; Figure 4-S4 shows the magnitude of cluster differences.). Although not shown here, the activation patterns across the four subgroups for contrasts of other emotional stimuli (i.e., happy, gruesome, and erotic) with neutral stimuli were similar to those shown above, and likewise, the general locations of the activation regions were similar for the four subgroups. Abbreviations: ALC_M_ = alcoholic men; ALC_W_ = alcoholic women; NC_M_ = nonalcoholic men; NC_W_ = nonalcoholic women.

**Figure 4-S4.**
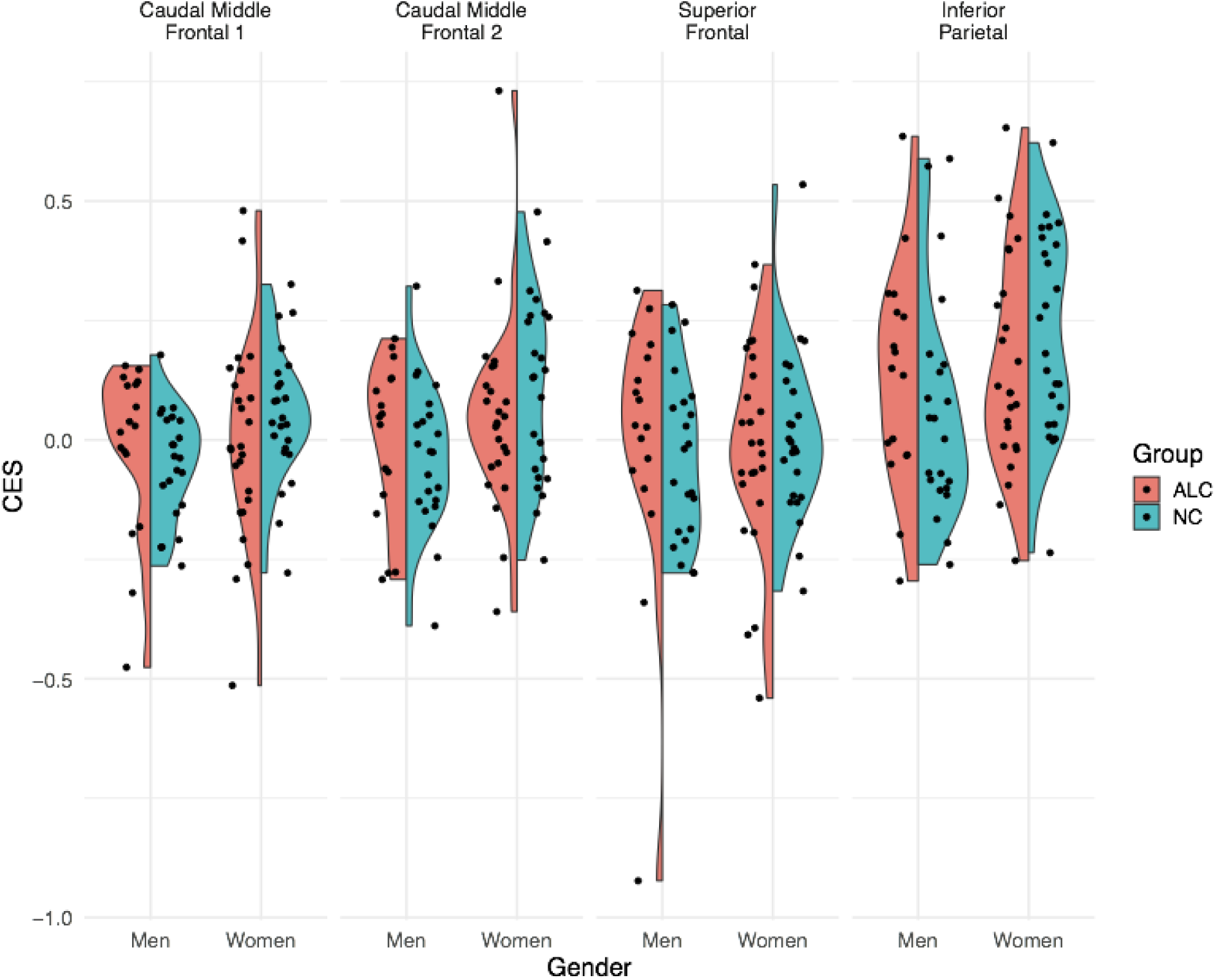
Contrast values observed in each cluster for aversive vs. neutral conditions. The split violin plot represents the Contrast Effect Size (CES; equivalent to “CON” in SPM or “COPE” in FSL) for left hemisphere clusters in which a significant group x gender interaction was identified for the aversive vs. neutral contrast ( *p*<0.05 after correction for multiple comparisons). Positive values indicate aversive > neutral, while negative values indicate aversive < neutral. Each point represents a single participant’s average CES for vertices within the cluster. This figure is meant to convey the variability in CES across participants that is not visible in Figure 4. In each of the four clusters, nonalcoholic control men had greater activation to aversive stimuli than neutral stimuli, and the contrast was more positive than was observed for alcoholic men. The pattern was reversed for women: alcoholic women had lower contrast values than nonalcoholic women. Abbreviations: ALC = alcoholic participants; NC = nonalcoholic participants.

**Table 2-S1.**
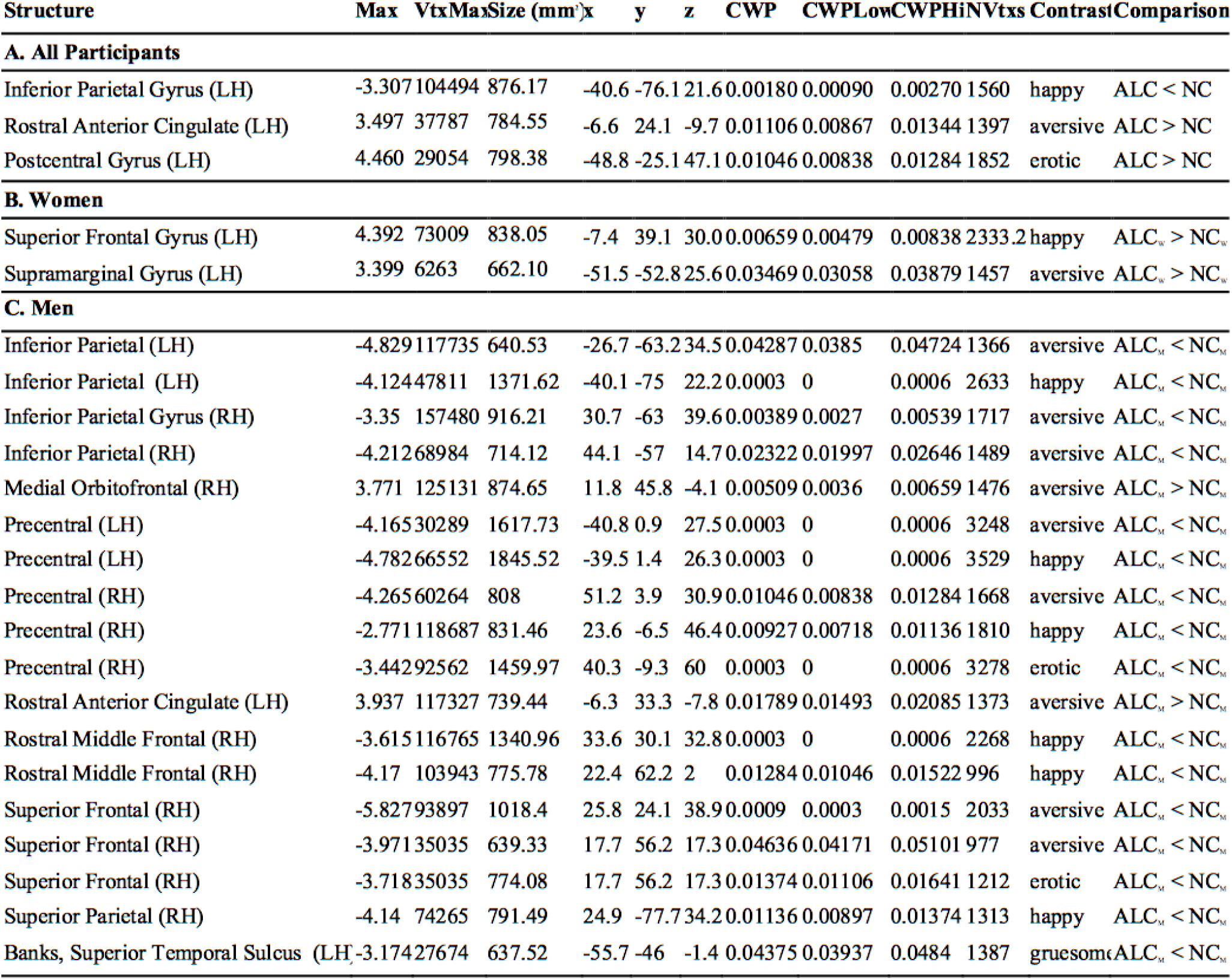
Cortical brain activation differences between alcoholics and nonalcoholic control subjects. MNI305 coordinates for peak voxel within significant clusters of activation showing difference between alcoholic and nonalcoholic control participants determined by surface-based whole brain analyses in (a) all subjects, (b) women only, and (c) men only. Abbreviations: LH = left hemisphere; RH = right hemisphere; Max = maximum −log10(*p*-value) in the cluster; VtxMax = vertex number at the maximum; size = surface area of cluster; XYZ = the MNI coordinates of the maximum; CWP = clusterwise *p-*value further corrected for the three spaces of left cortex, right cortex, and volume; CWP Low and CWPHi - 90% confidence interval for CWP; NVtxs = number of vertices in the cluster; ALC = alcoholic participants; NC = nonalcoholic Control participants.

**Table 2-S2.**
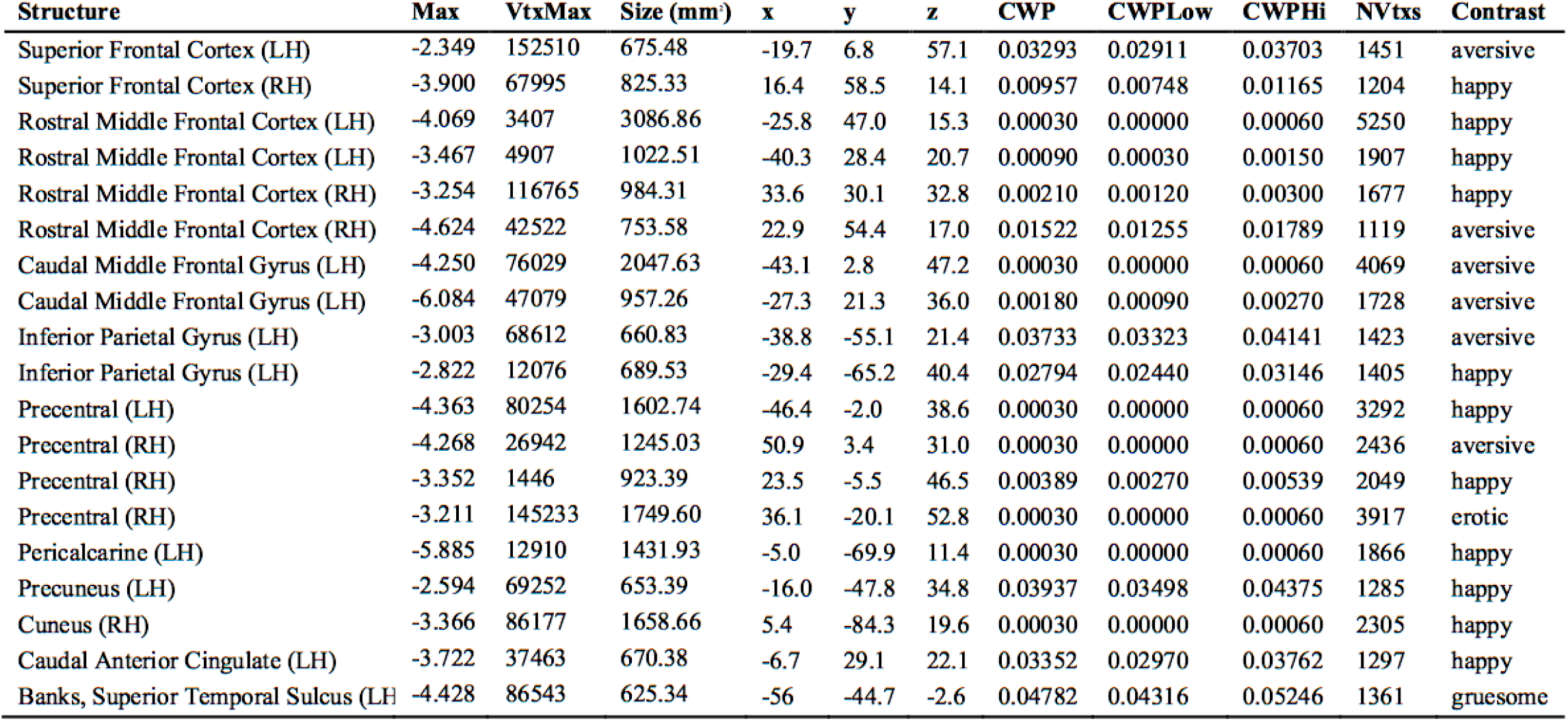
Cortical brain activation regions corresponding to the interactions between gender and alcoholism. MNI305 coordinates for peak voxel within significant clusters of activation showing group x gender interaction for emotion (happy, aversive, gruesome, and erotic vs. neutral) from surface-based, and volumetric whole brain analyses. Abbreviations: LH = left hemisphere; RH = right hemisphere; Max = maximum −log10(*p*-value) in the cluster; VtxMax = vertex number at the maximum; Size = surface area of cluster; XYZ = Montreal Neurological Institute (MNI) coordinates of the maximum; CWP = clusterwise *p*-value further corrected for the three spaces of left cortex, right cortex, and volume; CWPLow and CWPHi = 90% confidence interval for CWP; NVtxs = number of vertices in the cluster.

**Table 2-S3.** Significant brain activation differences determined through volumetric based comparisons. MNI305 coordinates for peak voxel within significant clusters of activation determined through volumetric whole brain analyses. Abbreviations: LH = left hemisphere; RH = right hemisphere; Max = maximum −log10(*p*-value) in the cluster; XYZ = Montreal Neurological Institute (MNI) coordinates of the maximum; CWP = clusterwise *p-*value further corrected for the three spaces of left cortex, right cortex, and volume.

| Structure | Size (mm) | x | y | z | CWP | Max | Comparison | Contrast |
| --- | --- | --- | --- | --- | --- | --- | --- | --- |
| Parahippocampal Cortex (LH) | 15920 | -34 | -23 | -27 | 0.0007912 | -5.62445 | Group x Gender | erotic |
| Accumbens (LH) | 50016 | -10 | 5 | -9 | 0.000000 | 5.49738 | Control: male vs. female | erotic |
| Cerebellum Cortex (LH) | 15960 | -14 | -39 | -23 | 0.00020 | -3.790 | Group x Gender | happy |
| Cerebellum Cortex (LH) | 7224 | -8 | -45 | -13 | 0.0467673 | 3.25617 | Control: male vs. female | happy |
| Thalamus (RH) | 12384 | 6 | -19 | 11 | 0.0016394 | -3.93561 | Male: alc vs. control | happy |
| Cerebellum Cortex (LH) | 18648 | -22 | -79 | -29 | 0.0000831 | 4.05785 | Control: male vs. female | aversive |
| Cerebellum Cortex (LH) | 15240 | -6 | -41 | -19 | 0.00049 | -3.409 | Group x Gender | aversive |

